# PKC modulator bryostatin-1 therapeutically targets CNS innate immunity to attenuate neuroinflammation and promote remyelination

**DOI:** 10.1101/2023.08.28.555084

**Authors:** Payam Gharibani, Efrat Abramson, Shruthi Shanmukha, Matthew D. Smith, Wesley H. Godfrey, Judy J. Lee, Jingwen Hu, Maryna Baydyuk, Marie-France Dorion, Xiaojing Deng, Yu Guo, Soonmyung Hwang, Jeffrey K. Huang, Peter A. Calabresi, Michael D. Kornberg, Paul M. Kim

**Author notes:** These authors contributed equally to this work (Payam Gharibani, Efrat Abramson).

## Abstract

In multiple sclerosis (MS), microglia and macrophages within the central nervous system (CNS) play an important role in determining the balance between myelin repair and demyelination/neurodegeneration. Phagocytic and regenerative functions of these CNS innate immune cells support remyelination, whereas chronic and maladaptive inflammatory activation promotes lesion expansion and disability, particularly in the progressive forms of MS. No currently approved drugs convincingly target microglia and macrophages within the CNS, contributing to the critical lack of therapies promoting remyelination and slowing progression in MS. Here, we found that the protein kinase C (PKC)-modulating drug bryostatin-1 (bryo-1), a CNS-penetrant compound with an established human safety profile, produces a shift in microglia and CNS macrophage transcriptional programs from pro-inflammatory to regenerative phenotypes, both in vitro and in vivo. Treatment of microglia with bryo-1 prevented the activation of neurotoxic astrocytes while stimulating scavenger pathways, phagocytosis, and secretion of factors that promote oligodendrocyte differentiation. In line with these findings, systemic treatment with bryo-1 augmented remyelination following a focal demyelinating injury in vivo. Our results demonstrate the potential of bryo-1 and functionally related PKC modulators as myelin regenerative and neuroprotective agents in MS and other neurologic diseases through therapeutic targeting of microglia and CNS-associated macrophages.

**One Sentence Summary:** PKC modulation in CNS innate immune cells favors the activation of a beneficial phenotype that promotes myelin regeneration and neuroprotection.

## INTRODUCTION

Multiple sclerosis (MS) is an inflammatory demyelinating disease of the central nervous system (CNS) that affects nearly three million people worldwide (*1*). MS most commonly begins with a relapsing course (termed relapsing-remitting MS or RRMS) defined by discrete episodes of focal CNS inflammation followed by variable, often incomplete recovery (*2, 3*). These attacks of inflammation are driven largely by the peripheral, adaptive immune system, characterized by dense lymphocytic infiltrates crossing the blood-brain barrier (BBB) and producing focal plaques of demyelination. Many individuals with an initial relapsing course ultimately convert to secondary progressive MS (SPMS), characterized by insidious worsening of disability in the absence of discrete relapses. A minority of individuals demonstrate a progressive course from onset, termed primary progressive MS (PPMS). Current disease-modifying therapies (DMTs) for MS primarily target peripheral, adaptive immunity, preventing relapses and new plaque formation. However, no approved therapies promote recovery of function through an induction of successful remyelination or slow the insidious worsening of disability that occurs independent of relapses in the progressive forms of MS (*4, 5*). The lack of therapies that promote remyelination and slow progression remains a critical unmet need.

Unlike RRMS, the pathology of progressive forms of MS is largely independent of peripheral inflammation, characterized instead by chronic activation of CNS innate immune cells (microglia and CNS-associated macrophages) both at the edges of chronic active plaques and diffusely throughout the CNS. Microglia/macrophages play dual roles in CNS injury and repair. Many chronic MS plaques contain a rim of activated, iron-laden microglia/macrophages, which are associated with lesion expansion, ongoing demyelination, and progressive disability (*6–10*). In animal models of inflammatory demyelination, these cells contribute to injury through a variety of maladaptive functions including antigen presentation, inflammatory cytokine production, release of cytotoxins, induction of toxic astrocytes, and complement-mediated synapse engulfment (*11–13*). Conversely, microglia/macrophages play critical roles in remyelination and repair through phagocytic clearance of inhibitory debris (e.g., myelin debris generated from demyelination), resolution of inflammation, and secretion of trophic factors that support neuroaxonal health and promote oligodendrocyte (OL) differentiation (*14, 15*). Therefore, a major goal of current drug discovery efforts is the identification of treatments that therapeutically modulate microglia and CNS-associated macrophages, thereby inhibiting maladaptive functions while promoting regenerative phenotypes (*5, 16*).

We previously reported that the protein kinase C (PKC)-modulating drug bryostatin-1 (bryo-1) potently attenuates neuroinflammation and neurologic disability in the experimental autoimmune encephalomyelitis (EAE) mouse model (*17*). In the periphery, bryo-1 selectively targeted innate immune cells (dendritic cells and peripheral macrophages) with limited direct actions on lymphocytes. In dendritic cells and macrophages, modulation of PKC with bryo-1 reciprocally inhibited inflammatory cytokine production (e.g., IL-6 and IL-12) while promoting an anti-inflammatory phenotype characterized by IL-10 and arginase-1 (Arg1) expression. Bryo-1 is a natural compound that binds to the C1 domain of PKC, which represents the conserved endogenous binding site for diacylglycerol (DAG) (*18*). DAG, bryo-1, and other PKC activating chemical compounds (such as phorbol esters) produce distinct downstream PKC functions upon binding to C1, owing to structurally distinct interactions of these molecules with PKC and the lipid bilayer (*19–21*). In this way, bryo-1 antagonizes the tumorigenic and mitogenic effects of phorbol esters, and thus, bryo-1 was initially tested as an anti-cancer drug (*22*). Intriguingly, we found that a subset of structurally unrelated PKC-binding agents (e.g., prostratin) produced effects on innate immunity similar to those observed with bryo-1, as did several synthetic analogs of bryo-1 (bryologs) (*23*). These findings identified bryo-1 and functionally related PKC modulators as potential therapeutic agents for targeting innate immune functions. Critically, bryo-1 has been investigated in human trials of cancer, HIV, and Alzheimer’s disease (AD) (*24–27*). While results have been mixed, it has demonstrated a favorable safety profile over up to two years of treatment. Bryo-1’s use in AD was based on preclinical findings that it induces neuronal synaptogenesis through a specific PKC isoform (PKCε) rather than any observed impact on immunity (*28–31*). As such, bryo-1 has only been investigated as a short-term, intermittent treatment intended to boost cognition rather than as a DMT altering AD progression (*25–27*).

Our previous work demonstrated that bryo-1, which has been shown to cross the BBB, produced neurologic improvement even in the late-stage EAE in wild-type C57BL/6 mice, when peripheral inflammation had subsided and smoldering neuroinflammation persists (*17*). Given the critical role of CNS innate immune cells in remyelination and progressive MS, we therefore wondered whether bryo-1 produces therapeutic effects on microglia/CNS macrophages similar to those observed in peripheral myeloid cells. In support of this possibility, PKC-signaling pathways have been implicated in key microglia functions (*32*). Perhaps most intriguingly, a gain-of-function mutation in phospholipase C gamma 2 (PLCγ2) lowers AD risk in humans (*33, 34*). PLCγ2 is predominantly expressed by microglia/CNS macrophages and is responsible for producing DAG, the endogenous PKC activator. Furthermore, PLCγ2 mediates signaling downstream of triggering receptor expressed on myeloid cells 2 (TREM2), a scavenger receptor necessary for CNS-resident macrophage protective functions, including phagocytosis and promotion of remyelination (*35–37*). A recent study also demonstrated that TREM2 impedes the complement-mediated synaptic pruning that occurs during neurodegeneration by forming a complex with complement component 1q (C1q) (*38*). Targeting PKC, which is further downstream of TREM2, may provide a unique advantage of a more specific functional regulation of microglia/CNS macrophage activity.

Here, we investigated the impact of PKC modulation with bryo-1 on microglia and CNS-associated macrophages both in vitro and in two distinct in vivo models of neuroinflammation and focal demyelination: EAE and lysolecithin (LPC)-induced demyelination, respectively. Bryo-1 treatment produced a transcriptional profile in these cells consistent with a regenerative phenotype, including downregulation of classically pro-inflammatory and neurotoxic genes along with upregulation of genes encoding trophic factors, scavenger receptors, and factors associated with resolution of inflammation. Consistent with these findings, bryo-1 limited the induction of neurotoxic astrocytes while promoting protective functions such as myelin phagocytosis and induction of oligodendrocyte precursor cell (OPC) differentiation. Most critically, we found that these beneficial effects of bryo-1 led to augmented OPC recruitment, differentiation, and remyelination following focal spinal cord demyelination in vivo. These findings solidify a role for PKC as a key regulator of microglia/macrophage function in the CNS, while identifying bryo-1 and functionally related PKC modulators as candidate drugs for promoting remyelination and limiting progression in MS and other neurologic diseases. Given its established safety profile, bryo-1 has a direct path to clinical development for these indications.

## RESULTS

### Bryo-1 modulates glial transcriptional responses to inflammatory stimuli

To examine the impact of bryo-1 in a system that preserves interactions between microglia and astrocytes, we used mixed glial cells (MGCs) derived from mouse cortices, as previously described (*23*). Because we previously found that bryo-1 modulates the response to pro-inflammatory signals in peripheral macrophages (*17*), we first examined bryo-1’s effects on transcriptional responses to lipopolysaccharide (LPS) stimulation (Fig. 1 and table S1). RNA sequencing (RNA-seq) demonstrated that co-treatment with bryo-1 substantially altered the response to LPS stimulation compared to vehicle co-treatment (Fig. 1A). Consistent with our findings in peripheral macrophages, Kyoto Encyclopedia of Genes and Genomes (KEGG) enrichment analysis found that bryo-1 downregulated many canonical pro-inflammatory pathways induced by LPS, including those associated with antigen processing and presentation; signaling pathways associated with toll-like receptors (TLRs), nucleotide oligomerization domain (NOD)-like receptors, nuclear factor kappa-light-chain-enhancer of activated B cells (NF-κB), and tumor necrosis factor (TNF); and the complement cascade (Fig. 1B). A similar overall effect of bryo-1 treatment was seen in MGCs co-stimulated with LPS and interferon-γ (IFNγ) (fig. S1A and table S2).

**Fig. 1.**
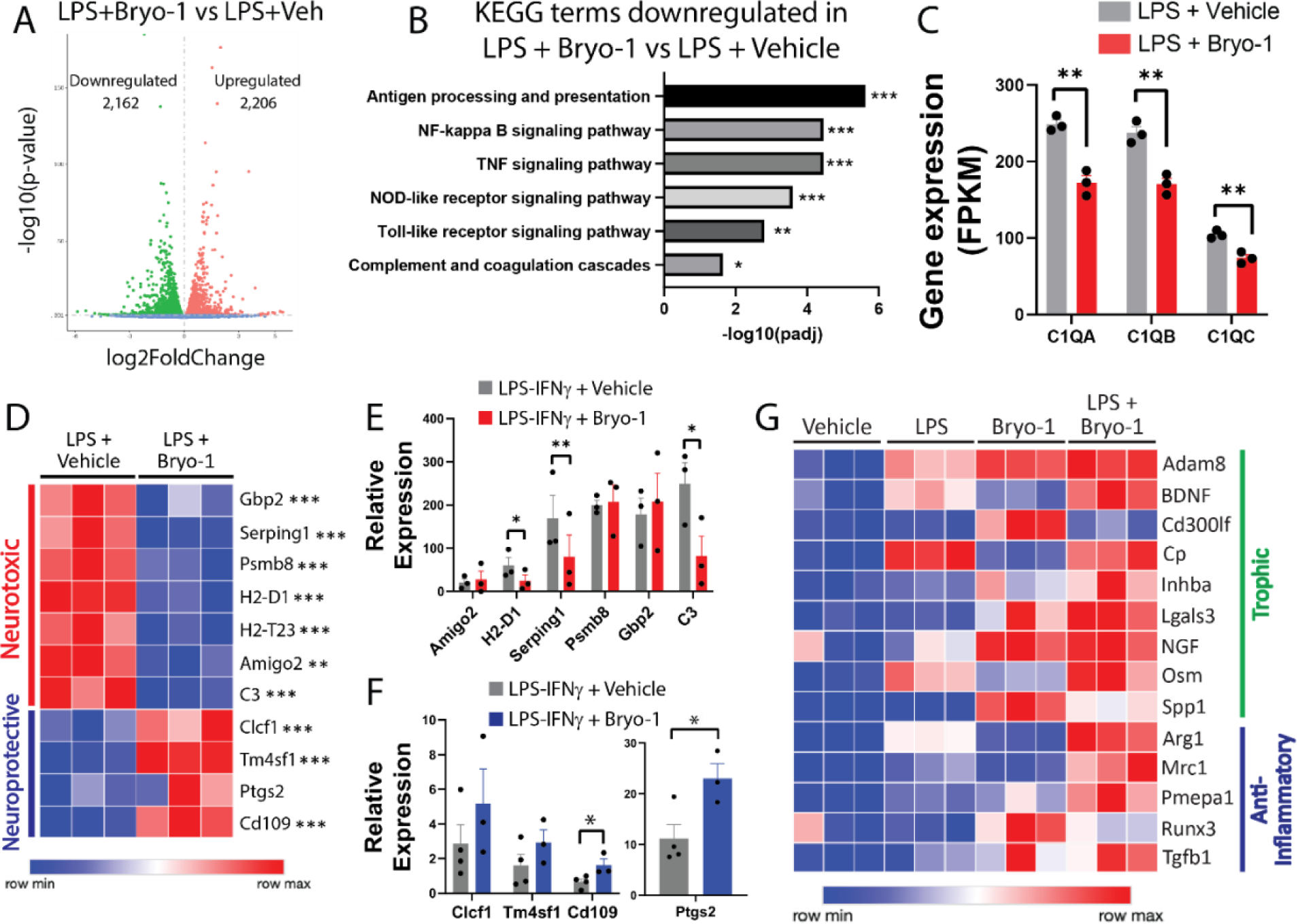
Bryo-1 modulates responses to inflammatory stimulation in cultured microglia and astrocytes. **(A-D)** Mouse cortical MGCs were treated for 24 hours with LPS (100 ng/ml) plus bryo-1 (50 nM) or vehicle, followed by RNA-seq analyses. **(A)** Volcano plot of genes up- and downregulated by bryo-1 in the presence of LPS. **(B)** KEGG enrichment analysis showing key inflammatory pathways downregulated by bryo-1 in LPS-stimulated glial cultures. **(C)** Bryo-1 decreases expression of C1q genes. **(D)** Bryo-1 modulates inflammatory astrocyte transcriptional programs induced by LPS in MGCs. **(E, F)** Effects of bryo-1 on astrocyte polarization are mediated by microglia. Microglia isolated from 4-week-old mouse brain were treated with LPS (100 ng/ml) and IFNγ (20 ng/ml) plus either bryo-1 (50 nM) or vehicle for 24 hours. Microglia-conditioned media was then added to primary astrocyte cultures for an additional 24 hours, followed by qPCR analysis from astrocytes for conventional neurotoxic **(E)** and neuroprotective **(F)** genes. Direct treatment of astrocytes with bryo-1, either in the presence of microglia-conditioned media or polarizing cocktail, had no effect on astrocyte transcriptional responses (fig. S2). **(G)** Bryo-1 augments expression of trophic and anti-inflammatory genes in cortical MGCs in the presence of LPS. The cells were prepared and stimulated as in A-D. Data derived from three biological replicates per condition (A-D, G) or represent mean ± SEM from 3-4 independent experiments (E, F). * p<0.05, ** p<0.01, *** p<0.001 using DESeq2 adjusted for false discovery rate (A-D, G) or two-tailed student’s t-test adjusted for multiple comparisons (E, F).

We were particularly intrigued by bryo-1-mediated regulation of the complement pathway, given the established role of complement signaling in microglia-astrocyte crosstalk and the pathophysiology of both MS and classic neurodegenerative diseases (*10, 13, 39–44*). In MGCs, bryo-1 significantly attenuated expression of the C1q subunits *C1QA*, *C1QB*, and *C1QC* (Fig. 1C). Suppression of the complement C1q subunits is significant because C1q is upregulated in microglia at the leading edge of chronic active MS lesions, termed microglia inflamed in MS (MIMS) (*10*), and is required for complement-mediated synaptic elimination and β-amyloid-mediated synaptic injury (*38, 41*).

To examine the impact of bryo-1 on astrocyte transcriptional profiles, we evaluated expression of genes associated with neurotoxic and neuroprotective astrocyte phenotypes (*40, 45*). Treatment with bryo-1 potently downregulated expression of neurotoxic astrocyte genes induced by stimulation with LPS (Fig. 1D) or LPS-IFNγ (fig. S1B), including complement *C3*. Under the same conditions, bryo-1 upregulated expression of astrocyte genes associated with a neuroprotective phenotype, e.g., *Clcf1*, *Tm4sf1*, *Ptgs2*, and *Cd109*.

Microglia and astrocytes were cultured separately to determine whether bryo-1’s effects on reactive astrocyte profiles are mediated by direct actions on astrocytes or indirectly through microglia modulation. As previously reported (*40*), the conditioned media from LPS-IFNγ-stimulated microglia potently induced expression of neurotoxic genes in cultured astrocytes (Fig. 1E). When microglia were co-treated with bryo-1, induction of neurotoxic astrocyte genes by the conditioned media from bryo-1-treated microglia was significantly attenuated (Fig. 1E), whereas expression of neuroprotective astrocyte genes was augmented (Fig. 1F). In contrast, direct treatment of astrocytes with bryo-1 had no effect on gene expression changes induced by the conditioned media from LPS-IFNγ-stimulated microglia (fig. S2A, B). Similarly, direct treatment of astrocytes did not alter neurotoxic gene expression induced by a cocktail of TNF, IL-1α, and C1q (fig. S2C). These findings indicate that bryo-1 directly modulates microglia’s inflammatory response, with downstream consequences for induction of reactive astrocytes.

Because bryo-1 upregulates anti-inflammatory cytokine expression in LPS-stimulated peripheral macrophages and dendritic cells (*17*), we examined bryo-1’s impact on expression of trophic and anti-inflammatory microglia genes in MGCs that were either unstimulated or stimulated with LPS (Fig. 1G) or LPS-IFNγ (fig. S3). Interestingly, bryo-1 augmented expression of several of these factors on its own but also acted synergistically with LPS and LPS-IFNγ to induce even greater expression of many such factors. Among the trophic factors induced by bryo-1 treatment were neurotrophins, such as *BDNF* and *NGF*, and genes associated with promotion of OL differentiation and remyelination, such as *Adam8*, *Cd300lf*, *Cp*, *Inhba* (activin A), *Lgals3* (galectin-3), *Osm* (oncostatin M), and *Spp1* (osteopontin) (*46–51*). Similarly, bryo-1 acted synergistically with LPS and LPS-IFNγ to induce genes associated with resolution of inflammation, such as *Arg1*, *Mrc1* (CD206), *Pmepa1*, *Runx3*, and *Tgfb1*.

Taken together, the above findings suggest that, analogous to its effects on peripheral innate immune cells, bryo-1 modulates inflammatory activation of microglia – downregulating classic inflammatory pathways and augmenting transcriptional programs associated with tissue repair and resolution of inflammation.

### Bryo-1 potentiates regenerative microglia transcriptional programs

We previously found that bryo-1 potentiates the effects of the typical type 2 inflammatory response on markers of a regenerative phenotype in peripheral macrophages (*17*). Based on this finding and the above results demonstrating that bryo-1 augments trophic and anti-inflammatory gene expression with LPS stimulation, we wanted to further explore bryo-1’s effects on microglia expression of pathways and specific genes associated with remyelination and tissue repair, with and without IL-4 stimulation. Using RNA-seq analyses, we found that treatment of unstimulated MGCs with bryo-1 produced substantial changes in gene expression (Fig. 2A and table S3). KEGG enrichment analysis revealed that bryo-1 upregulated key pathways associated with protective/regenerative microglia functions, including pathways associated with phagocytosis, proliferation, and cell motility (Fig. 2B). As with peripheral macrophages, these effects were further potentiated in the presence of IL-4, such that combined treatment with bryo-1 and IL-4 upregulated these pathways to a greater extent than treatment with either bryo-1 or IL-4 alone (Fig. 2C, D and table S4).

**Fig. 2.**
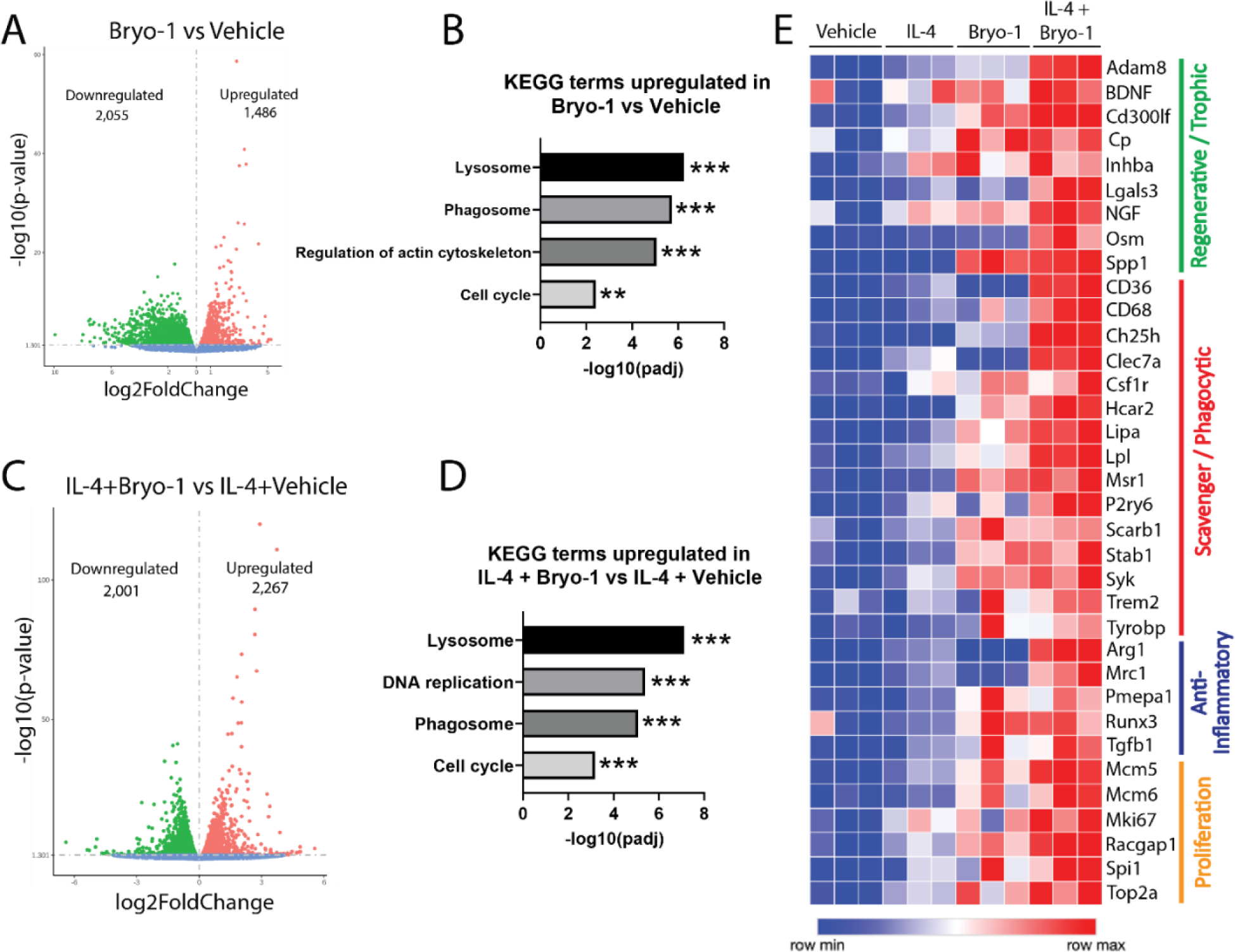
Bryo-1 promotes a regenerative transcriptional profile in MGCs. Mouse cortical MGCs were treated for 24 hours with bryo-1 (50 nM) or vehicle ± IL-4 (20 ng/ml), followed by RNA-seq analyses. **(A)** Volcano plot of genes up- and downregulated by bryo-1 compared to vehicle. **(B)** KEGG enrichment analysis showing key phagocytic and proliferative pathways upregulated by bryo-1. **(C)** Volcano plot of genes up- and downregulated by IL-4 plus bryo-1 compared to IL-4 plus vehicle. **(D)** KEGG enrichment analysis showing key phagocytic and proliferative pathways upregulated by IL-4 plus bryo-1 compared to IL-4 plus vehicle. **(E)** Heat map demonstrating that bryo-1 upregulates a wide range of trophic, phagocytic, anti-inflammatory, and proliferative genes associated with regenerative microglia functions, both alone and synergistically with IL-4. Data derived from three biological replicates per condition. * p<0.05, ** p<0.01, *** p<0.001 using DESeq2 adjusted for false discovery rate.

To better understand the impact of bryo-1 on microglia phenotype, we examined bryo-1 regulation of specific microglia genes reported to play key roles in regenerative and protective functions (Fig. 2E). As reported above, bryo-1 treatment alone increased expression of many genes encoding trophic factors and proteins associated with resolution of inflammation. Similar to peripheral macrophages, expression of these genes was substantially potentiated by combined stimulation with bryo-1 and IL-4. In the case of Arg1, protein expression in the presence of IL-4 was dramatically increased by treatment with as little as 1 nM bryo-1 (fig. S4A), with expression exclusively within CD11b^+^ cells (fig. S4B). Microglia proliferation is critical to tissue repair and protection (*14, 15*), and in line with our KEGG analysis, bryo-1 upregulated many microglia genes associated with a proliferative state. Finally, Bryo-1, alone and/or synergistically with IL-4, substantially increased the expression of specific scavenger receptors and related genes reported to promote remyelination and neuroprotection through phagocytosis of myelin debris and neurotoxic plaques (*14, 15*). Among the scavenger-related genes upregulated by bryo-1 were *CD36*, *Clec7a*, and multiple components of TREM2 signaling, including *TREM2*, its associated adaptor gene/protein *Tyrobp*/DAP12, and downstream kinase *SYK* (*52, 53*). These findings demonstrate that bryo-1 promotes regenerative/protective microglia transcriptional programs, which is further potentiated in the setting of endogenous pro-regenerative and anti-inflammatory signals.

### Functional analyses of bryo-1-treated microglia and CNS-derived macrophages

We next examined whether the bryo-1-mediated changes in gene transcription identified by our RNA-seq experiments translated to functional modulation of microglia and CNS-derived macrophages. Microglia/macrophage phagocytic function is critical for remyelination in order to clear myelin debris that serves as a potent inhibitor of OL differentiation. To determine the effect of bryo-1 on phagocytosis by microglia, we first used primary mouse microglia cultures incubated with fluorescent-tagged myelin. Treatment with bryo-1 significantly increased phagocytic uptake of myelin (Fig. 3A). To determine whether systemically administered bryo-1 sufficiently enters the intact CNS and augments phagocytosis, we used an ex vivo phagocytosis assay. Mice were treated with bryo-1 or vehicle daily for three days via intraperitoneal (IP) injection. On day 4, the dissociated cells from the brains were incubated with fluorescent-tagged myelin for four hours followed by an evaluation of myelin engulfment by microglia/CNS macrophages via flow cytometry. We found that systemic treatment with bryo-1 significantly increased the myelin-phagocytic capacity of microglia and CNS macrophages (Fig. 3B-D; gating strategy shown in fig. S5).

**Fig. 3.**
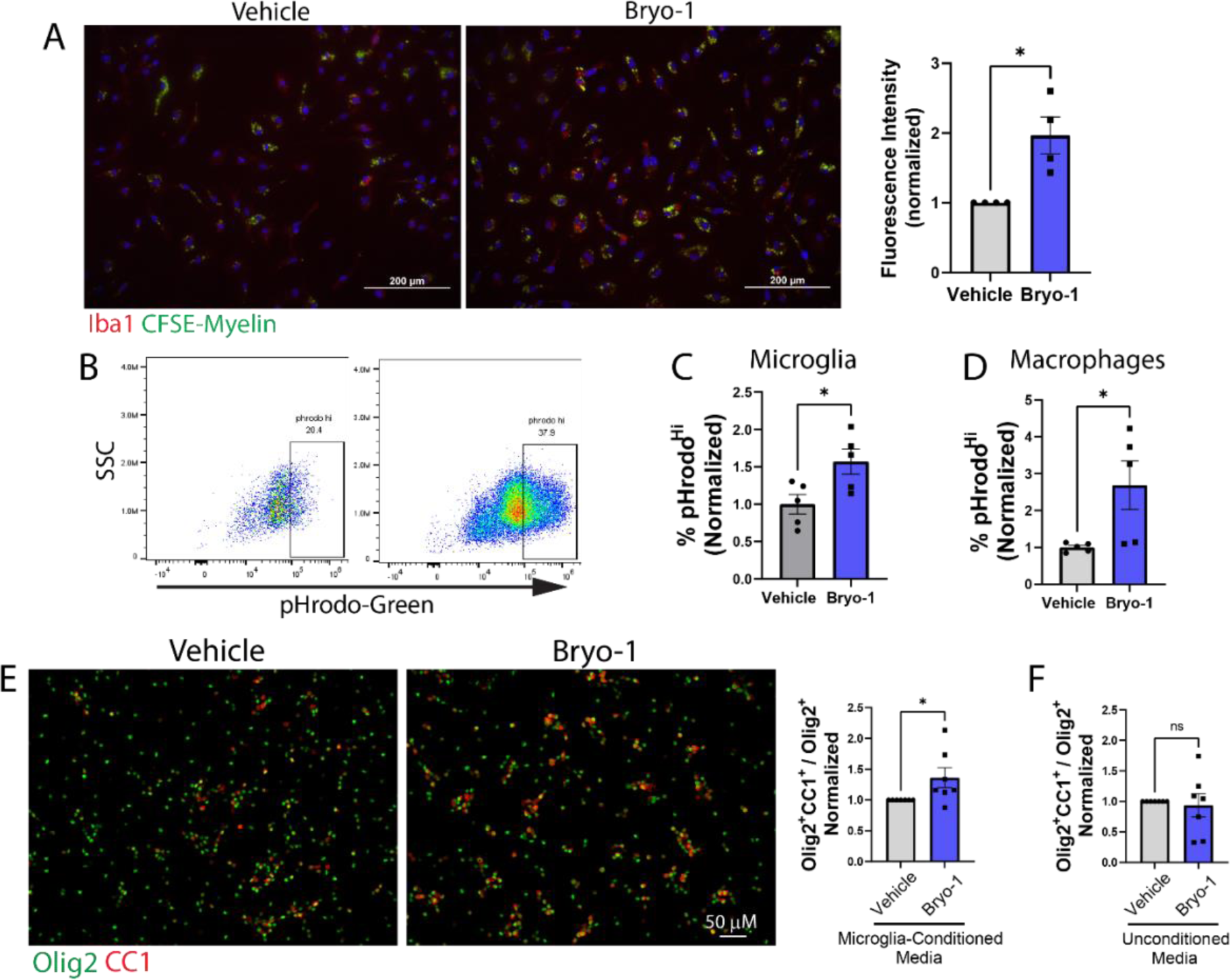
Bryo-1-treated microglia and brain macrophages demonstrate enhanced phagocytosis and induction of OPC differentiation. **(A)** Primary mouse microglia were treated with bryo-1 (50 nM) or vehicle for 48 hours followed by 2-hour incubation with CFSE-labeled myelin. Phagocytosis was measured as CSFE fluorescence intensity within Iba1^+^ cells. Representative images (*left*) and mean ± SEM from n=4 independent experiments (*right*). **(B**-**D)** Mice were treated with bryo-1 (35 nmol/kg, a dose chosen based on our previous work) (*17, 23*) or vehicle daily for 3 days by IP injection. On day 4, brains were dissociated and incubated with pHrodo-tagged myelin for 4 hours, followed by flow cytometric analysis. CD45^+^CD11b^+^ cells were identified as microglia or macrophages based on Clec12a expression. Representative flow plot **(B)** and mean ± SEM from n=5 mice per group **(C, D)**. Gating strategies are shown in figure S5. **(E)** Conditioned media derived from primary mouse microglia that were treated with bryo-1 (50 nM) or vehicle for 48 hours was collected and applied to cultured mouse OPCs for 48 hours. OPC differentiation was then quantified by IF staining for CC1. Representative images (*left*) and mean ± SEM from n=7 independent experiments (*right*). **(F)** To examine direct effects of bryo-1 on OPCs, bryo-1 (50 nM) or vehicle was added to microglia media (in the absence of microglia) for 48 hours, and the unconditioned media was then applied to cultured mouse OPCs for an additional 48 hours prior to quantification of OPC differentiation as in (E). Data represent mean ± SEM from n=7 independent experiments. ns = not significant, * p<0.05 by two-tailed student’s t-test.

Another critical regenerative function of microglia is the secretion of trophic factors that promote OL differentiation. Two key microglia-secreted proteins known to induce OL differentiation are activin A (*46*) and galectin-3 (*48*), which are encoded by *Inhba* and *Lgals3* genes, respectively. Our RNA-seq analyses demonstrated upregulation of these genes by bryo-1 (Figs. 1G, 2E, and S3). As such, we reasoned that bryo-1 might enhance the ability of microglia to induce OL differentiation. To test this hypothesis, primary mouse microglia were treated with bryo-1 or vehicle for 48 hours. The microglia-conditioned media was then applied to cultured OPCs for 48 hours, and OL differentiation was assayed by immunofluorescent (IF) staining. As hypothesized, OL differentiation was increased when OPCs were cultured with the conditioned media from bryo-1-treated microglia (Fig. 3E). To rule out direct actions of bryo-1 on OPCs, media was treated identically with bryo-1 or vehicle for 48 hours in the absence of microglia, and this unconditioned media was then applied to OPCs for an additional 48 hours. The bryo-1-unconditioned media had no effect on OL differentiation (Fig. 3F).

### Bryo-1 modulates inflammatory and regenerative microglia and CNS-associated macrophage responses in vivo

We next sought further evidence of whether the in vitro and ex vivo effects of bryo-1 on microglia/CNS macrophages similarly occur with systemic bryo-1 treatment in vivo. To this end, we used two mouse models to examine microglia/macrophage behavior in distinct environments: 1) myelin oligodendrocyte glycoprotein peptide 35-55 (MOG_35-55_)/complete Freund’s adjuvant (CFA) EAE, which is characterized predominantly by neuroinflammation, and 2) LPC-induced focal demyelination, which allows examination of microglia/macrophage activity in the context of demyelination and remyelination.

We previously found that systemic treatment with bryo-1 (35 nmol/kg) via IP injection three times per week potently attenuated neurologic deficits in C57BL/6 mice subjected to MOG_35-55_/CFA EAE, even when treatment began during or past the peak of disease (*17, 23*). To best isolate bryo-1’s effects on microglia/macrophages in the EAE model, we began treatment with bryo-1 or vehicle at post-immunization day (PID) 19, representing late-peak disease when the acute, adaptive inflammatory response has subsided and chronic neuroinflammation by microglia/macrophages persists. Mice were treated with bryo-1 or vehicle three days per week until PID 35 (representing the chronic stage of the model), at which point microglia/CNS macrophages were isolated from brain and spinal cord via fluorescence-activated cell sorting (FACS) and evaluated by RNA-seq analyses (schematic shown in Fig. 4A). Similar to our findings in MGCs exposed to inflammatory stimuli, systemic treatment with bryo-1 downregulated classic pro-inflammatory pathways in microglia/CNS macrophages compared to vehicle treatment while augmenting pathways associated with regeneration/repair and resolution of inflammation (Fig. 4B-D and table S5).

**Fig. 4.**
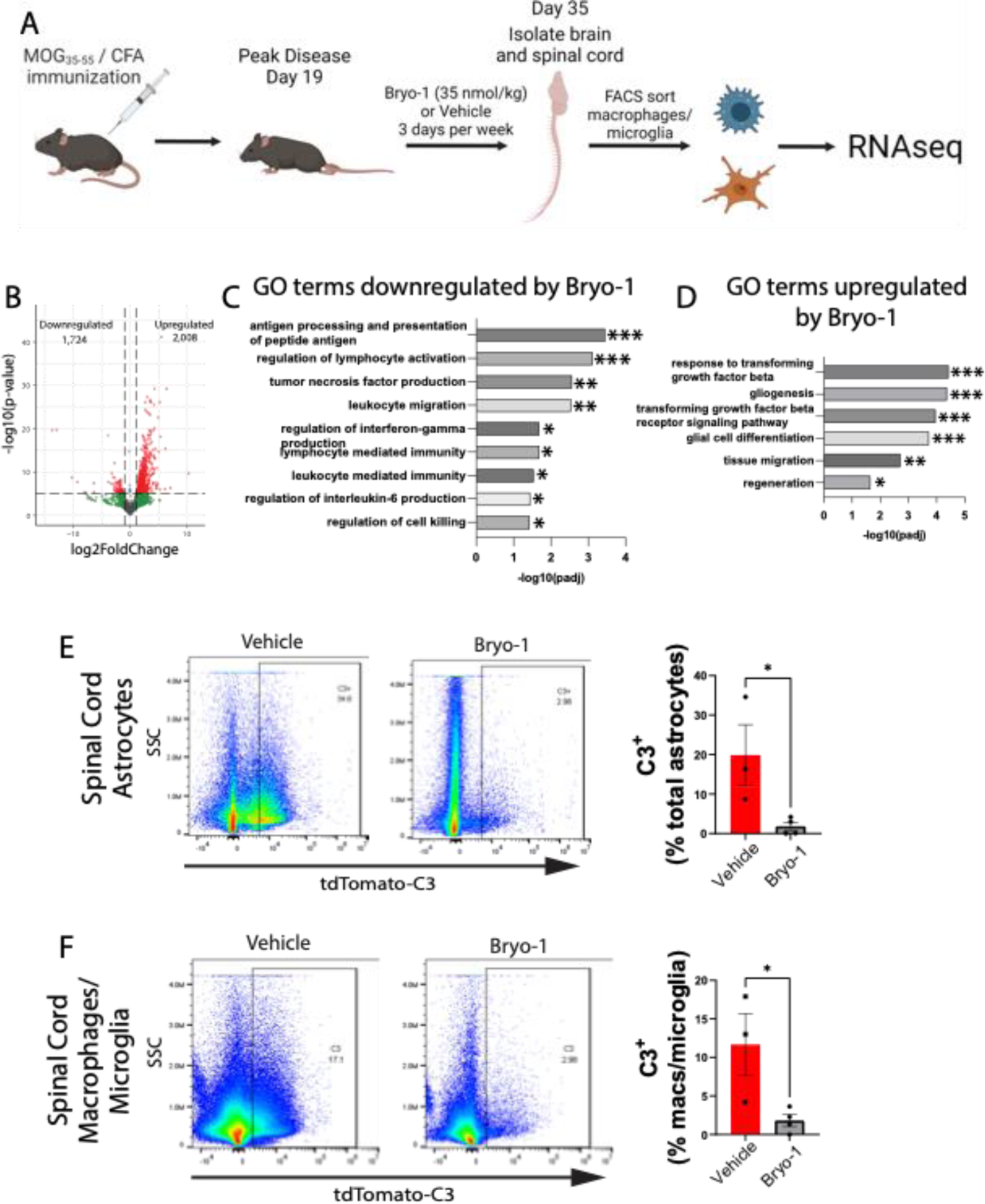
Bryo-1 therapeutically modulates CNS innate immune cell phenotypes during inflammatory demyelination in vivo. **(A-D)** Mice subjected to EAE were treated with bryo-1 or vehicle beginning on PID 19. On PID 35, microglia/macrophages were flow sorted from the CNS, followed by RNA-seq analyses. **(A)** Schematic of experimental design. **(B)** Volcano plot of genes up- and downregulated by bryo-1 compared to vehicle. Bryo-1 downregulated pro-inflammatory gene programs **(C)** and upregulated regenerative programs **(D)**, similar to the effects observed in MGCs. n=3 mice per group. **(E, F)** Mice subjected to EAE were treated with bryo-1 or vehicle beginning at the onset of neurologic symptoms. On PID 19, mice were sacrificed and C3 expression in astrocytes **(E)** and microglia/macrophages **(F)** was analyzed by flow cytometry from the spinal cord. Representative flow plot (*left*) and mean ± SEM from n=3 mice per group (*right*). Gating strategies are shown in figure S6. * p<0.05, ** p<0.01, *** p<0.001 using DESeq2 adjusted for false discovery rate (C, D) or two-tailed student’s t-test (E, F).

To supplement our RNA-seq analyses and evaluate microglia-astrocyte crosstalk, we used C3-tdTomato reporter mice (*42, 54*) to examine the impact of bryo-1 on C3 expression under neuroinflammatory conditions. In these conditions, C3 is expressed mostly by astrocytes and to a lesser extent by microglia/CNS macrophages. C3 expression is a marker of neurotoxic reactive astrocytes, which are induced by inflammatory microglia (*40*). C3 directly contributes to the neuronal loss in both inflammatory and neurodegenerative contexts (*13, 41, 42, 44*), and C3 genetic variants have been linked to faster rates of neurodegeneration in MS (*43*). Mice were again subjected to MOG_35-55_/CFA EAE. Bryo-1 or vehicle treatment was initiated at the onset of neurologic deficits, and C3 expression was evaluated by flow cytometry on PID 19. Bryo-1 treatment potently suppressed C3 expression in both astrocytes (Fig. 4E) and microglia/CNS macrophages (Fig. 4F), in agreement with our in vitro findings (gating strategy shown in fig. S6).

The above findings demonstrate that bryo-1 favorably modulates microglia/CNS macrophage responses to inflammation in vivo. However, remyelination is limited in EAE owing to substantial axonal injury that occurs in this model. To explore the impact of bryo-1 on CNS innate immune responses during remyelination, we induced focal demyelination in mouse ventral spinal cord via stereotaxic injection of LPC. To avoid effects of drug treatment on initial OL injury, treatment with bryo-1 was initiated at 2 days post-lesion (dpl), when demyelination is complete (schematic of experimental design shown in Fig. 5A). Lesions were examined at early (9 dpl) and late (15 dpl) periods of OL differentiation and remyelination. It was previously shown that successful remyelination requires the transition from a population of iNOS^+^ inflammatory microglia to an Arg1-expressing regenerative population (*46, 47*). We observed similar results in our model, with an initial wave of iNOS^+^/Iba1^+^ cells being replaced by Arg1^+^/Iba1^+^ cells (Fig. 5B-D). Bryo-1 treatment promoted this transition, decreasing the population of iNOS^+^ cells at 9 dpl (Fig. 5C) and increasing the population of Arg1^+^ cells at 15 dpl (Fig. 5D). Quantitative PCR (qPCR) performed from the lesions demonstrated that bryo-1 treatment increased expression of many scavenger/phagocytic genes with roles in myelin debris clearance and remyelination (Fig. 5E), corroborating our in vitro findings.

**Fig. 5.**
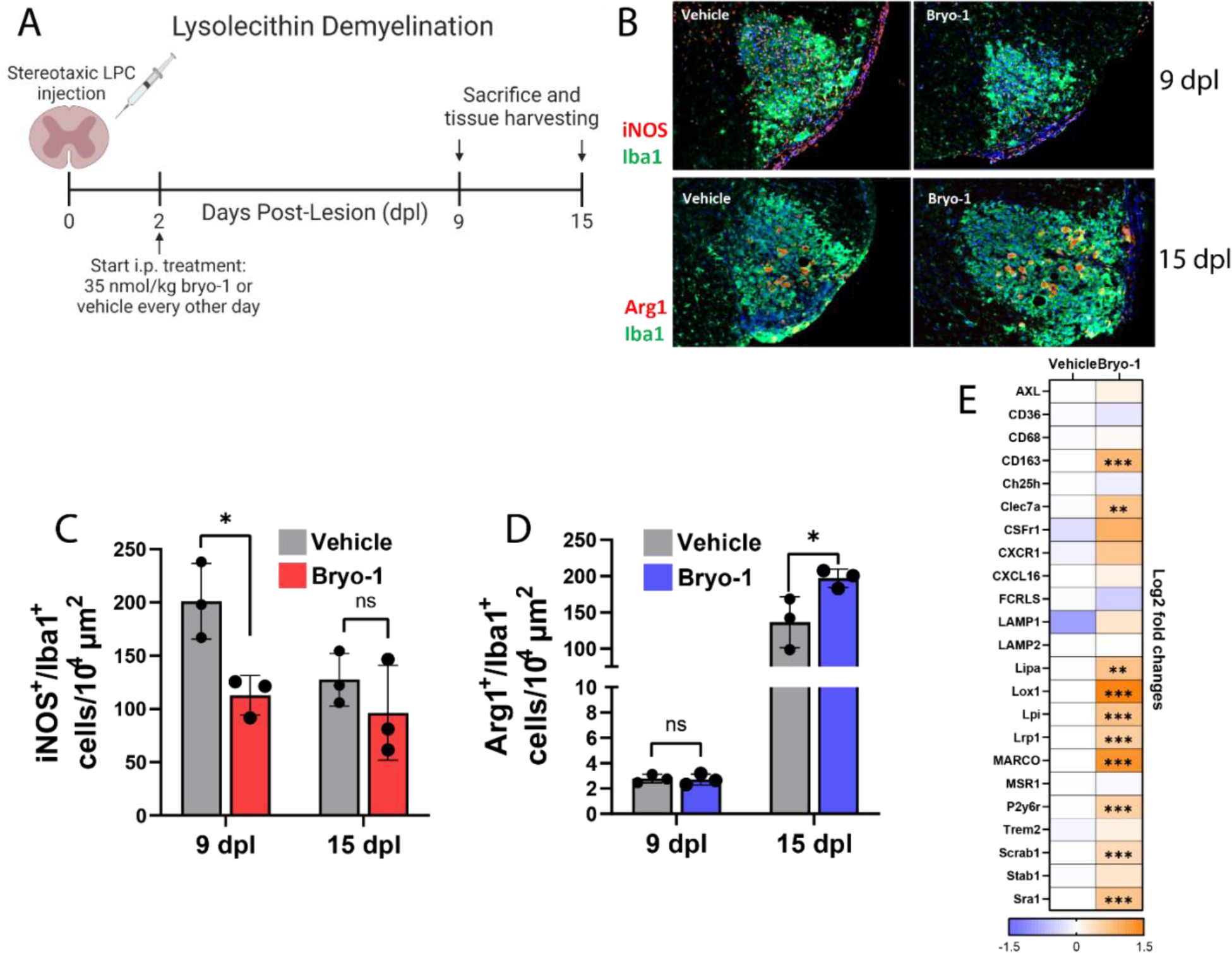
Systemic treatment with bryo-1 promotes a regenerative innate immune response following focal CNS demyelination. **(A)** Schematic of experimental design. Focal demyelination was induced in the ventral column of the spinal cord. **(B-D)** Arg1 and iNOS expression were quantified in Iba1^+^ microglia/macrophages within the lesions by IF staining at 9 and 15 dpl. Representative image **(B)** of iNOS (*top row*) and Arg1 (*bottom row*) staining at 9 and 15 dpl, respectively, and quantification of iNOS **(C)** and Arg1 **(D)** expression. Data represent mean ± SEM from n=3 mice per group. **(E)** qPCR analysis from focally demyelinated lesions at 9 dpl demonstrates that bryo-1 treatment increases expression of scavenger receptors with known roles in myelin debris clearance and remyelination. Data derived from n=3 mice per group. * p<0.05, ** p<0.01, *** p<0.001 by two-tailed student’s t-test adjusted for multiple comparisons.

Taken together, the above findings show that systemic treatment with bryo-1 effectively targets microglia and CNS-associated macrophages in vivo, limiting maladaptive inflammatory responses and favoring responses associated with remyelination and repair.

### Systemic treatment with bryo-1 augments remyelination in vivo

To determine whether the modulation of microglia/CNS macrophages described above translates into therapeutic enhancement of remyelination, we again used the LPC-induced focal demyelination model, with bryo-1 or vehicle treatment beginning on 2 dpl. Systemic treatment with bryo-1 led to increased numbers of OL-lineage cells and mature OLs within lesions at both 9 and 15 dpl (Fig. 6A, B). Consistent with these findings, qPCR performed from the lesioned spinal cord tissue demonstrated increased expression of *Olig2* as well as genes encoding the myelin proteins, myelin basic protein (MBP) and myelin-associated glycoprotein (MAG) (Fig. 6C). To evaluate remyelination directly, we performed IF staining for MBP and transmission electron microscopy (TEM) for measurement of g-ratios at 15 dpl. Expression of MBP was increased within the lesions in bryo-1-treated mice (Fig. 6D). TEM analysis revealed more myelinated axons and decreased g-ratios in mice treated with bryo-1 (Fig. 6E-G), demonstrating the augmentation of successful remyelination by bryo-1.

**Fig. 6.**
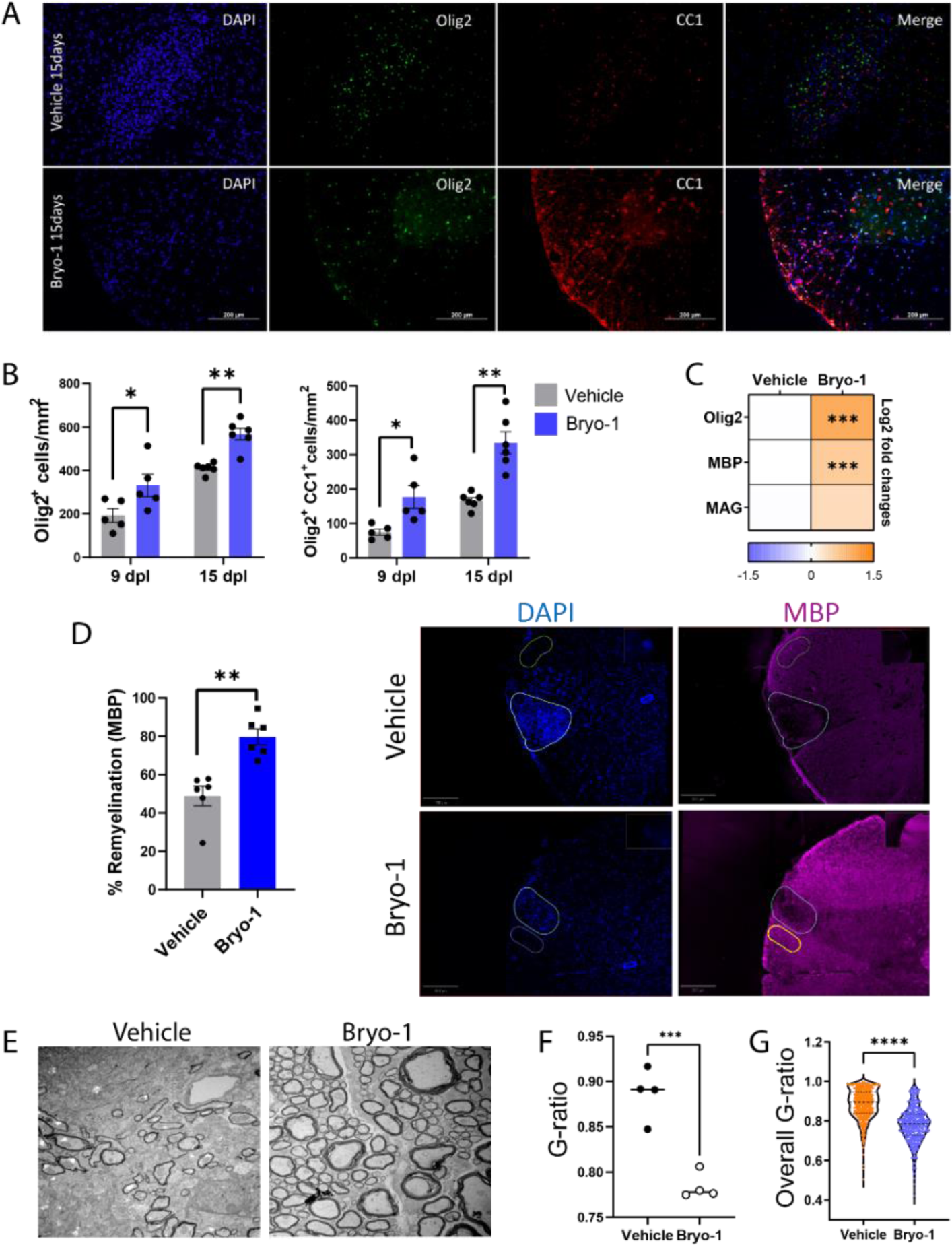
Bryo-1 enhances OPC differentiation and remyelination following focal demyelination. Experimental design was as summarized in Fig. 5A. **(A)** Representative images of lesions from 15 dpl in bryo-1 and vehicle treated mice. **(B)** Quantification of total OL-lineage cells (*left*, Olig2^+^) and differentiating OL (*right*, Olig2^+^CC1^+^) within lesions at 9 and 15 dpl. Data represent mean ± SEM from 5-6 mice/group. (**C**) qPCR of myelin-specific gene expression within lesions at 9 dpl. Data derived from 3 mice/group. (**D**) Remyelination within lesions at 15 dpl, as measured by MBP expression. MBP expression was measured in an automated fashion, based on fluorescence intensity within the lesion compared to an adjacent non-lesioned site. Quantification (mean ± SEM) from 6 mice/group (*left*) and representative images (*right*) with lesioned and non-lesioned sites outlined. **(E)** Representative TEM images of myelinated axons. **(F)** Quantification of g-ratios from n=4 mice/group, with a minimum of 5 fields/mouse. **(G)** Quantification pooled from all measured axons (436 for vehicle and 365 for bryo-1). * p<0.05, ** p<0.01, *** p<0.001, **** p<0.0001 by two-tailed student’s t-test, adjusted for multiple comparisons where appropriate.

MS is a female-predominant disease, occurring with approximately 3:1 (female:male) ratio (*55*). Furthermore, sex differences have been reported in CNS innate immune functions and remyelination (*56*). For these reasons, we examined the effects of bryo-1 treatment as a function of sex, finding that bryo-1 increased total OL-lineage cells, OL differentiation, and remyelination in both female and male mice (fig. S7).

## DISCUSSION

In this study, we found that PKC modulation by bryo-1, a CNS-penetrant natural compound with an established human safety profile, therapeutically regulates microglia/CNS macrophages. This distinct modulation limits maladaptive, neurotoxic functions while augmenting activities associated with neuroprotection and regeneration, thereby enhancing remyelination in vivo. Given the critical role of microglia and CNS-associated macrophages in progressive MS and other neurodegenerative diseases that lack effective therapies, identifying treatments that manipulate these cells in a clinically favorable fashion remains a vital but currently unmet clinical goal. Our findings identify PKC as a relevant target for future studies. Most importantly, these results identify bryo-1 as a lead candidate drug for modulating microglia and CNS-associated macrophages with a low threshold for clinical translation based on existing human clinical trial experience for other conditions.

Our prior work identified structurally unrelated PKC-binding drugs, as well as synthetically derived bryologs, that share the beneficial effects of bryo-1 in cultured peripheral myeloid cells and the EAE model (*23*). Although investigations of these agents in the models used for our current work was beyond the scope of the study, it seems likely that many of these compounds will similarly replicate the effects of bryo-1 in microglia/CNS macrophages. These additional PKC-modulating agents represent a chemical library and pipeline for identifying drugs with potential clinical advantages, such as improved tolerance, a wider therapeutic window, and ease of synthesis and formulation. Future work will focus on developing screens for identifying promising candidates within this pipeline.

Although our work on bryo-1 focused on remyelination and neuroinflammation due to its effects on CNS innate immune cells, the biological rationale underlying recent bryo-1 clinical studies in AD was based on mouse studies demonstrating that bryo-1 acutely induces neuronal synaptogenesis through PKCε. Informed by this biological mechanism, previous phase 2 studies of bryo-1 in AD sought to identify *improvements* in cognition after a short course of bryo-1 treatment administered every other week. These studies utilized an ambitious clinical design, treating patients with moderate-to-severe AD for 3-6 months with a primary outcome of improvement in the Severe Impairment Battery cognitive scale by weeks 13 or 28. After initially mixed results, the most recent study was reported to have missed its primary outcome (*25, 27*). Based on this result, we anticipate that bryo-1’s application as a microglia/CNS macrophage-targeting agent for neuroinflammatory and neurodegenerative disorders will require a longer treatment duration with more frequent dosing than the previous AD trials to detect clinical benefit. Similarly, future studies should focus on long-term repair and slowed progression rather than short-term effects, such as synaptic plasticity. Similar to the lesson of anti-amyloid therapies, bryo-1’s benefit will likely be greatest for those treated early in the disease course, prior to extensive neurodegeneration.

Relatedly, there are important caveats to further clinical evaluation of bryo-1 in MS and other neuroinflammatory conditions. To date, bryo-1 has only been administered to humans intravenously, with doses given at most weekly. Although target engagement of PKC has been studied in the periphery and CNS penetrance has been established in animals, the optimal dosing regimen for CNS indications in humans has not been established. Before proceeding with large studies utilizing clinical outcomes, it will be critical to perform further dose-finding studies to establish optimized dosing for CNS penetrance and target engagement in people. Biomarkers of microglia activation, such as PET radioligands (*57, 58*) and MRI phase rim lesions (*6*), should aid in these studies.

Some additional limitations merit discussion. Although we implicate PKC as a therapeutic target in microglia and CNS-associated macrophages based on the actions of bryo-1, we included no genetic evidence of its role in this study, nor did we entirely exclude the possibility that off-target effects of bryo-1 mediate its actions. Nonetheless, the role of PKC isoforms as the mediators of bryo-1 actions on innate immunity is supported by our prior work showing that a bryolog with abrogated PKC binding failed to replicate bryo-1’s benefit in the EAE model and cultured myeloid cells (*23*). From a practical perspective, pharmacologic effects on PKC predict the functional consequences of bryo-1 derivatives and structurally unrelated PKC modulators (*19, 21, 59–61*). However, future studies should investigate the key signaling pathways up- and downstream of PKC that mediate bryo-1’s beneficial effects. Such studies will provide further biological insight into the physiologic regulation of microglia/CNS macrophages while identifying additional therapeutic targets. Previous reports of the therapeutic role of PLCγ2 (*35*), which acts upstream of PKC and downstream of TREM2, raises the intriguing possibility that bryo-1 and other PKC modulators might rescue deficient TREM2 activation – a possibility currently under investigation.

Extrapolation of our findings to progressive MS is also limited by the lack of animal models that sufficiently replicate the pathology of progressive MS. All EAE models involve an element of peripherally derived inflammation, such that differentiating drug effects in the periphery versus the CNS is challenging. To limit the influence of peripheral, adaptive immunity, our work has focused on later stages of C57BL/6 EAE, after the peak of peripheral immunity. We previously found that bryo-1 leads to improvement of neurologic functions even when treatment begins in the chronic phase of EAE, i.e., at PID 28 (*17*). In our current study, we investigated bryo-1’s effects on microglia/CNS macrophage profiles only after peak disease. Although no faithful model for progressive MS exists, the potential of bryo-1 as a therapeutic agent targeting microglia and CNS-associated macrophages in these patients is underscored by its benefit in a second model of demyelination, LPC-induced focal demyelination.

## MATERIALS AND METHODS

### Study design

The objective of the study was to determine the effects of PKC modulation with bryo-1 on CNS innate immunity and how these effects impact neuroinflammation and remyelination. Based on our prior work demonstrating that bryo-1 and functionally related PKC modulators favorably altered the phenotype of peripheral myeloid cells, the starting hypothesis was that bryo-1 would have similar effects on CNS-derived macrophages and microglia, although at study onset we planned to evaluate direct effects on astrocytes and OPCs/OLs as well. To accomplish the study objective, we used primary cell cultures derived from mouse brain (MGCs, microglia, astrocytes, and OPCs) as well as in vivo mouse models, EAE and LPC-mediated focal demyelination. Assay measurements/endpoints were all prospectively selected. Experimental design and the sample sets from which data were derived are described in detail in the main text, figure legends, and below. For in vivo studies, mice were randomized into treatment groups, with randomization and treatment performed by different investigators. Analyses were performed by investigators blinded to treatment group.

### Statistical analyses

For RNA-seq, differential expression testing was done with DESeq2 (*62*). Genes with adjusted p-value less than 0.05 were considered differentially expressed. KEGG and gene ontology (GO) pathway enrichment on the differentially expressed genes was performed with clusterProfiler (*63*), and terms with adjusted p-values less than 0.05 were considered significantly enriched. For all other studies, statistics were performed using GraphPad Prism 9.0. Statistical significance was determined using two-tailed student’s t-tests, adjusted for multiple comparisons where appropriate. The number of mice, independent experiments, and technical or biological replicates from which the data were derived are described in the figure legends.

### Mice

C57BL/6J mice were purchased from The Jackson Laboratory (stock # 000664). CD1 IGS mice were purchased from Charles River (strain code 022). C3-tdTomato reporter mice were previously described (*54*). All mice were housed in a pathogen-free, temperature-controlled Johns Hopkins (JH) mouse facility. All protocols were approved by the JH Institutional Animal Care and Use Committee.

### Bryo-1 preparation/treatments

Bryo-1 was purchased from Tocris Bioscience (catalog no. 2383). For treatment of cultured cells, bryo-1 was dissolved in 100% ethanol to a final working concentration of 50 mM, which was stored at 4°C for up to 3 months. The working bryo-1 solution was added to cultured cells to yield the final drug concentrations indicated in the manuscript, with 100% ethanol serving as vehicle control. For mouse treatment, bryo-1 was diluted to a concentration of 7 nmol/ml in 20% ethanol in PBS, which was stored at -20°C until use. Mice were treated via IP injection with 35 nmol/kg bryo-1 or vehicle control (20% ethanol in PBS).

### Primary cell cultures

#### MGCs

MGCs were prepared from P3-6 C57BL/6 mouse as described (*23*). Cortices were isolated and placed in a 15 ml centrifuge tube with Hanks’ Balanced Salt Solution (HBSS). Excess buffer was removed, and 1 ml of 0.05% trypsin was added per brain. The cortices were dissociated in trypsin with a 10 ml serological pipet and incubated for 15-20 min at 37°C in a water bath. Trypsin was neutralized with equal volume of Dulbecco’s Modified Eagle Media (DMEM)/F12, containing 1% antibiotics and 10% fetal bovine serum (FBS). The dissociated cortices were triturated with 6 ml HBSS with a 5-ml serological pipet fixed to a 1-ml sterile pipet tip, filtered twice through a 100-µm cell strainer into a 50-ml conical tube, and centrifuged at 1,000 rpm for 10 min. The cell pellet was resuspended in DMEM/F12 media and plated on 6-well plates (250,000 cells/ml), pre-coated with 100 mg/ml poly-D-lysine (PDL). The media was replaced every three days until confluency on 14 days in vitro (DIV).

#### Microglia

Microglia were isolated from 4-week-old CD1 mice as previously described (*64*). In brief, the cerebra were cut into fine pieces in a cold enzyme digestion buffer containing HBSS (without calcium/magnesium), 5% FBS, 10 μM HEPES, 2 mg/ml collagenase D, and 28 U/ml DNase I. A single-cell suspension was obtained by trituration. This suspension was filtered through a 70-µM filter and centrifuged 10 min at 300 g at 4°C, followed by a debris removal procedure (*64*). The obtained cell pellet was allowed to bind to CD11b magnetic microbeads (Miltenyi Biotec) for 15 min on ice with gentle mixing every 5 min, followed by magnetic separation of CD11b^+^ cells using a MACS LS column (Miltenyi Biotec). The cells obtained in CD11b^+^ fractions were centrifuged and resuspended in DMEM/F12 supplemented with 1% penicillin-streptomycin, 2 mM glutamine, 5 µg/ml N-acetyl-L-cysteine, 1× SATO [100 µg/ml human apo-transferrin (Sigma), 100 µg/ml BSA, 16 µg/ml putrescine (Sigma), 60 ng/ml progesterone (Sigma; from stock: 2.5 mg /100 μl ethanol), 40 ng/ml sodium selenite (4.0 mg/100 μl 0.1 N NaOH in neurobasal medium)], 10% FBS, and 20 ng/ml M-CSF. Microglia were maintained in culture for 7 DIV before use in downstream assays.

#### Astrocytes

Forebrain and midbrain from P1-P7 C57BL6/J mice were isolated and placed in cold HBSS without cations. The tissues were dissociated on ice and spun at 300 g for 2 min. The mixture was then resuspended in Enzyme P using Neural Tissue Dissociation Kit P (Miltenyi Biotec, catalog no. 130-092-628) and was spun at 37°C for 15 min, followed by Enzyme A addition. The mixture was triturated 15-20 times with a serological pipette, followed by a 10-min incubation. This step was repeated twice with successively smaller pipette tips to achieve thorough dissociation. HBSS with cations were added to the mixture. The cells were passed through a 70-μm filter and, following centrifugation, resuspended in MACS buffer and Fc block. ACSA2 beads were added according to the manufacturer’s protocol recommended concentration. After the bead incubation, cells were spun and resuspended for LS column depletion (Miltenyi Biotec, catalog no. 130-042-401). The cells were then added to the LS columns and washed three times with MACS buffer. The column was removed from the magnet, and 5 ml of MACS buffer was used to elute the cells with a plunger. The cells were spun down, resuspended in 1× SATO, and counted for plating (*65*). Astrocytes were maintained in culture for 7 DIV before use.

#### OPCs

Cerebral cortices from P4-6 CD1 mouse pups were dissected in ice-cold HBSS without Mg^2+^ and Ca^2+^ and triturated with 2 ml of digestion buffer containing 20 U/ml papain (Worthington) and 100 U/ml DNase I (Worthington) into single-cell suspension. Digestion was stopped by adding HBSS with Mg^2+^ and Ca^2+^. The cells were filtered through a 100-μm filter, and OPCs were isolated through immunopanning as described (*66*). Endothelial cells and microglia were depleted with BSL-1 and CD11b panning plates. OPCs were collected with PDGFRa panning plate, and 20,000 cells were seeded onto glass cover slips coated with PDL. Seeded OPCs were kept in proliferating media containing 40 ng/ml PDGF-AA (Peprotech), 10 ng/ml CNTF (Peprotech), and 1 ng/ml NT3 (Peprotech) for 48 hours before use.

### Astrocyte polarization

#### Astrocyte polarization with microglia-conditioned media

Microglia were treated with 100 ng/ml LPS and 20 ng/ml INFγ for 24 hours. Microglia-conditioned media was then collected and centrifuged to remove dead cells. Protein concentrations of media were measured and normalized to 50 μg/ml. The conditioned media was then added with protease inhibitors to astrocytes. Following 24 hours, astrocytes were harvested on ice and prepared using the universal protocol of qPCR with SYBR Green. Microglia were treated with 50 nM bryo-1 or vehicle (ethanol) at the time of stimulation with LPS-IFNγ. In a separate set of experiments, astrocytes were treated directly with 50 nM bryo-1 or vehicle at the same time as the microglia-conditioned media addition.

#### Astrocyte polarization with A1 cocktail

Astrocytes at 7 DIV were treated for 24 hours with 30 ng/ml TNF, 3 ng/ml IL-1α, and 400 ng/ml C1q, along with either 50 nM bryo-1 or vehicle.

### OPC differentiation

The microglia were cultured on PDL-treated 24-well plate at 2.5×10^4^ cells/well and treated with 50 nM bryo-1 or vehicle for 48 hours. The media was collected and centrifuged at 1500 rpm for 5 min to avoid any cell contamination in the conditioned media. OPC base media was mixed with microglia-conditioned media at a ratio of 1:1. Only CNTF was added with no PDGF-AA or NT3. After OPCs were treated for 48 hours with the conditioned media, OPCs were fixed with 4% PFA and permeabilized with 0.1% Triton X-100. The cells were blocked with 3% BSA/PBS for 1 hour and incubated with rabbit anti-Olig2 (1:500, Millipore), mouse anti-CC1 (1:100, Milllipore), and rat anti-MBP (1:50, Millipore) diluted in 1% BSA/PBS overnight. The cells were further stained with fluorescent secondary antibodies and DAPI. Slides were imaged on an epifluorescence microscope at four random fields. Differentiation analysis was done using Fiji software where Olig2^+^ and Olig2^+^CC1^+^ cells were counted in a blinded fashion.

### Myelin isolation/labeling

#### Myelin isolation

Myelin isolation and labeling with carboxyfluorescein succinimidyl ester (CFSE) were performed as described (*67*). Briefly, brains were dissected from ten 8-10 weeks old C57BL/6J mice and placed in ice-cold 0.32 M sucrose solution. The brains were cut into small pieces (∼5 mm^3^), transferred to 30 ml ice-cold 0.32 M sucrose solution, and then grounded into a smooth homogenate with a sterile hand-held rotary homogenizer. The brain homogenate was then gently transferred onto the top of ice-cold 0.83 M sucrose solution within a 50-ml polypropylene centrifuge tube, forming a sucrose density gradient and centrifuged at 100,000 g at 4°C for 45 min in a pre-cooled ultracentrifuge rotor. Myelin debris (white band) was collected from the interface between the two sucrose densities, homogenized in a sterile hand-held rotary homogenizer for 30-60 sec, and then centrifuged at 100,000 g at 4°C for 45 min. The solid white myelin pellet was resuspended in Tris-Cl solution (pH 7.4) and then centrifuged at 100,000 g at 4°C for 45 min. The pellet was suspended in sterile PBS and centrifuged at 22,000 × g for 10 min at 4 °C. The myelin pellet was resuspended in PBS to 100 mg/ml concentration.

#### Myelin labeling

Myelin (100 µl) from the above isolation protocol was resuspended in 200 µl of 50 µM CFSE or pHrodo-Green (Thermo Fisher Scientific), incubated at room temperature (RT) for 30 min in the dark, and centrifuged at 14,800 × g for 10 min at 4°C. The pellet containing labeled myelin was then washed three times in 100 mM glycine in PBS and resuspended in sterile PBS to a final concentration of 100 mg/ml. For pHrodo-Green labeling, 18.5 μl myelin was mixed with 25 μl pHrodo-Green and resuspended in 206.5 µl PBS (pH 8). The mixture was incubated in a RT rocker in the dark for 45 min. Myelin mixture was then centrifuged at 4,000 g for 10 min, supernatant was discarded, and resuspended in PBS (pH 7.2).

### In vitro microglia myelin phagocytosis

Microglia were cultured on PDL-treated glass coverslip in a 24-well cell culture plate at 2.5×10^4^ cells per well. These cells were treated 48 hours with 50 nM bryo-1 or vehicle, incubated with 1 mg/ml of CFSE-labeled myelin for 2 hours at 37°C, and washed three times with PBS to remove non-engulfed myelin debris. Microglia were fixed with 4% paraformaldehyde (PFA), blocked in 10% mouse serum, and incubated overnight at 4°C with Iba1 primary anti-goat antibody (1:250). This was followed by washing and incubation with a secondary antibody conjugated with Alexa Fluor 555 (1:500) for 45 min at RT and DAPI. The images were captured with a fluorescent microscope, and the fluorescence intensity within the microglia was calculated using ImageJ software. Iba1^+^ cells, which were completely in frame and did not overlap with other cells, were traced using ImageJ’s free hand selection tool. After each traced cell was added to the ROI manager, the highlighted Iba1^+^ ROI manager was superimposed into CFSE-labeled myelin channel. The exported data measured the mean fluorescence intensity of phagocytosed myelin for each microglia selected.

### Tissue preparation for phagocytosis/FACS

Brains and spinal cords from mice were collected into PBS on ice with 5 μg/ml actinomycin and 10 μM triptolide (*68*) and mechanically dissociated through a 16G needle attached to a 3-ml syringe five times. The dissociated samples were spun for 2 min at 200 g and resuspended in papain digestion buffer. The samples were incubated for 10 min at 37°C with slow rotation, followed by trituration with a p1000 pipet tip 10 times. Incubation and trituration were repeated, then passed over a 100-μm filter. The samples were spun for 10 min and resuspended in PBS and Miltenyi debris removal solution to remove myelin following manufacturer’s protocol.

### Ex vivo myelin phagocytosis

C57BL/6 mice (8-10 weeks old) were treated with 35 nmol/kg bryo-1 on days 0, 2, and 3 via IP injection. On day 4, the brains were prepared as described above, and the phagocytosis assay was performed as described (*69*). The dissociated cells were resuspended in DMEM/F-12 containing 10% (v/v) FBS in a conical tube, and pHrodo-conjugated myelin (3 μg) was added. The cells were incubated for 4 hours at 37°C, washed sequentially with PBS and PBS containing 2% FBS and 2.5 mM EDTA, and transferred to a 96-well conical-bottom plate. The cells were incubated with TruStain Fc and Zombie NIR (1:1500) for 20 min at RT. The cells were washed, stained with anti-CD11b BV510 (1:400; clone M1/70; Biolegend), anti-CD45 BV605 (1:400; clone 30-11; Biolegend), anti-Ly6G (1:400; clone 1A8; Biolegend), and anti-Clec12a APC (1:400; clone 5D3; Biolegend) antibodies for 30 min at RT, washed twice, and analyzed by flow cytometry on a Cytek Aurora Spectral flow cytometer.

### EAE

EAE was induced in 8-12 weeks old female C57BL/6J mice as previously described (*17, 70*). MOG_35-55_ peptide (amino acid sequence MEVGWYRSPFSRVVHLYRNGK) was prepared by the JH Synthesis and Sequencing Core Facility. CFA was prepared by adding 8 mg/ml of heat– killed Mycobacterium tuberculosis H37 Ra (Difco, catalog no. 231141) to incomplete Freund’s adjuvant (Thermo Fisher Scientific, catalog no. 77145). MOG_35-55_ peptide (2 mg/ml in PBS) was mixed 1:1 with CFA to make an emulsion. On day 0, mice were immunized by injecting 50 μl of the emulsion subcutaneously (SC) into each of two sites on the lateral abdomen. On days 0 and 2, the mice were injected IP with 250 ng pertussis toxin (List Biologicals, catalog no. 181) dissolved in PBS. On day 7, they were weighed and began scoring.

### Flow cytometry/flow sorting

Following brain/spinal cord dissociation and myelin removal from C57BL/6 or C3-tdTomato reporter mice, the cells were counted and incubated with 1 ml of TruStain Fc (1:200) and Live/Dead Aqua (1:2,000) for 20 min at RT. The cells were spun and resuspended in 300 μl surface stain (0.5% BSA, 1mM EDTA) in addition to these markers: Ly6G eFluor450 (1:250), CD11b PerCP-Cy5.5 (1:250), CD45 APCFire750 (1:250), and ACSA2 VioBright515 (1:50). The cells were stained for 30 min at RT, washed in 1 ml of 0.5% BSA and 1 mM EDTA, resuspended in 200 μl of 0.5% BSA, and filtered over a strainer FACS tube. The cells were analyzed either by flow cytometry with a Cytek Auorora or taken to JH Flow Cytometry Shared Resource Center for FACS sorting.

### RNA-seq

For all experiments, RNA was isolated using the Qiagen RNeasy Mini Plus Kit. For RNA-seq from MGCs, library preparation, sequencing, and analysis were performed by NovoGene. For RNA-seq of ex vivo flow-sorted myeloid cells, library preparation and sequencing were performed by the JH Genetic Resources Core Facility.

#### MGCs

Raw fastq files were processed with fastp (*71*), and reads containing adapter poly-N sequences or reads with low quality data were removed. Cleaned fastq files were then mapped to reference genome GRCm38 (mm10) with STAR (v2.6.1.d, parameter mismatch = 2) (*71*). FeatureCounts (v1.5.0p3) (*72*) was used to count the number of reads mapped to each gene, and reads per kilobase million (RPKM) was calculated based on length of the gene. Differential expression testing was done with DESeq2 (v1.20.0) (*62*). KEGG pathway enrichment on the up- and down-regulated differentially expressed genes was performed with clusterProfiler (*63*).

#### Flow-sorted myeloid cells

Raw fastq files were quantified using Salmon (1.4.0) (*73*) against a decoy aware RefSeq GRCm39 (*74*) transcriptome generated by concatenating the GRCm39 genome to the GRCm39 transcriptome. Transcripts were then summarized to gene level counts using tximport (1.26.1) (*75*). Differential expression testing was performed with DESeq2 (1.38.3). To avoid inflated fold changes resulting from low expression genes, log2 fold changes were shrunk with apeglm (*76*). GO term (*77, 78*) enrichment in the differentially expressed genes was performed using clusterProfiler (4.6.2). Volcano plot was constructed using EnhancedVolcano (1.16.0) (https://doi.org/10.18129/B9.bioc.EnhancedVolcano).

### Focal spinal cord demyelination

#### Demyelination

We induced focal demyelination by injecting 1% LPC (Sigma) in PBS into the ventral funiculus of the spinal cord of 8-12 weeks old C57BL/6J mice as described previously (*79*). Briefly, mice were anesthetized, and under a surgical microscope and on a stereotactic frame, a small midline incision was made below the ears on the back of the animal in the caudal direction. After visualizing T2, the spinal cord was exposed by a shallow cut in the connective tissue and ligamentum flavum between T3 and T4, followed by removing the dura with a 32G metal needle. Using a 34G needle (Hamilton Co.) connected to a 10 µl Hamilton syringe, 0.5 µl of 1% LPC was microinjected stereotaxically into the right ventral funiculus (depth of 1.3 mm) at 0.25 μl/min using a microinjection syringe pump (UMP3; World Precision Instrument). The needle was maintained in the spinal cord for 2 min after each injection to prevent efflux. Muscle and adipose tissues were closed with a single absorbable suture (Vicryl 5-0). Skin incision was closed with rodent wound clips. Saline (1 ml) and buprenorphine SR (1 mg/kg; ZooPharm, LLC) was administered after the surgery. Gentamycin (2 mg/kg; Henry Schein Animal Health) was administered SC every 12 hours for 3 days. On 2 dpl, treatment with 35 nmol/kg bryo-1 or vehicle was started by IP injection every other day. For the remyelination study, mice received IP injection of 500 μl of 1% neutral red (NR; Sigma-Aldrich) in PBS for spinal cord lesion labeling 2 hours prior to intracardiac perfusion with ice-cold 4% PFA for IF staining (*80*). For studying the innate immune cells, a separate cohort of LPC-injected mice was without NR staining. Bryo-1’s effects on the innate immunity and remyelination were studied by RT-qPCR at 9 dpl (female; n=5) and IF at 9 (female; n=5) and 15 dpl (female; n=3). To validate the remyelination effects of bryo-1 in both sexes, we also utilized male mice (n=3) for IF at 15 dpl.

#### Spinal cord IF staining

The spinal cords from mice perfused with 4% PFA were harvested and sectioned into a 3-mm piece with the epicenter (site of injection) of the lesion site in the middle. The tissues were post-fixed in 4% PFA at 4°C overnight. The tissues were incubated in gradient sucrose (10% then 30% sucrose in PBS) overnight for cryoprotection and embedded in OCT. The spinal cords were coronally sectioned into 12-µm sections at 0.8 mm from the epicenter at both sides (rostrally and caudally), collected on Superfrost Plus slides (VWR International), and dried for 30 min at RT before storing at -80°C. To reduce the NR intensity of the sections pertaining to remyelination studies, the samples were de-stained with a solution of 50% ethanol/1% glacial acetic acid for 10 min before performing IF. The sections were retrieved by Antigen Retrieval Reagent-Universal (R&D Systems) in a steamer for 10 min at 100°C, permeabilized with 1% Triton X-100 in TBS for 5 min, and incubated in blocking solution (10% donkey serum, 0.25% Triton X-100 in TBS) for 1 hour at RT. Primary antibodies were diluted in the blocking solution and applied overnight at 4°C. For remyelination studies, triple staining was carried out with rabbit anti-Olig2 (1:100, Millipore), mouse anti-CC1 (1:100, Sigma), and rat anti-MBP (1:50, Millipore). For studying CNS inmate immune cells, the tissues were stained with goat anti-Iba1 (1:100, Novus Biological), either with rabbit anti-Arg1 (1:100, Cell Signaling) or mouse anti-iNOS (1:200, Santa Cruz). After the primary antibodies, the sections were incubated with appropriate secondary antibodies (1:2,000) and counterstained with DAPI. A minimum of six sections (three rostral and three caudal) were counted manually by investigators who were blinded to the conditions. The total numbers of positive cells were normalized to the area of the injury using ImageJ. MBP staining was quantified by measuring the pixel intensity segmentation at the lesion site (normalized to a non-lesion site) using the open source QuPath software (https://qupath.github.io) (*81*). Images were taken by Zeiss Axio Observer Z1 epifluorescence microscope at 20x magnification (Zeiss).

#### RT-qPCR

Total RNA was isolated and reverse transcribed into cDNA with QuantiTect Reverse Transcription Kit. RT-qPCR was performed in triplicates for each sample using PowerUp SYBR Green Master Mix (Applied Biosystems) with a CFX384 Real-Time System (Bio-Rad). 10 ng of cDNA was used for each reaction. Primers are listed in table S6. Relative expression values were calculated using the 2^−ΔΔCT^ method and normalized against peptidylprolyl isomerase-A (ppia) or β-2-microglobulin (B2m) for spinal cord samples or GAPDH for astrocytes.

#### TEM

Mice (female; n=4) at 15 dpl were perfused with PFA prepared in 0.1 M Sorensen’s buffer (Electron Microscopy Sciences). The spinal cords were harvested, and 3 mm of lesion sites were cut with the epicenter in the middle. The specimens were incubated in 0.1 M Sorensen’s buffer (pH 7.2) containing 4% PFA and 3% glutaraldehyde for 1 hour at RT, before placing them at 4°C for 48 hours. The fixation buffer was replaced with 0.1 M Sorenson’s buffer. The blocks corresponding to the area of interest (∼0.8 mm from the epicenter) were trimmed and post-fixed in 2% osmium tetroxide (Electron Microscopy Sciences). The tissues were then dehydrated in graded ethanol and embedded in Embed 812 (EMS). Semi-thin sections were stained with toluidine blue for validating the area. Thin sections (70-80 nm) were cut on a Reichert Jung Ultracut E microtome and placed on copper 200 mesh grids. The grids were stained with uranyl acetate, which was followed by lead citrate. All images were captured on a Hitachi TEM at 80 kV. Total percentage of remyelinated axons were averaged from 5 images/group/mouse (area = ∼400 µm^2^/image). The g-ratio was calculated as the ratio of the inner to outer diameter of myelinated axons; a minimum of 100 fibers per mouse was analyzed (365 for bryo-1 and 436 for vehicle). The investigators involved in this process were blinded to the conditions.

## Acknowledgments

The authors thank L. Albacarys, J. P. Antel, L. M. Healy, S. H. Snyder, and M. Wang for their advice and support. We are also very grateful to C. Kemper (NHLBI, NIH), who developed and provided the C3 reporter mice to P. Calabresi. Cytek Aurora Spectral flow cytometer used in C3 reporter mouse experiments is funded by an NIH Grant (S10OD026859).

## Funding

Department of Defense, MS200232 (MDK, PMK)

TEDCO Maryland Innovative Initiative, 135025 (MDK, PMK)

Race to Erase MS, Young Investigator Award (MDK)

NIH MSTP Grant T32 GM136577 (WHG)

American Association of Immunologists Careers in Immunology Fellowship (MDK, WHG)

National Institutes of Health, R01NS041435 (PAC)

National Institutes of Health, 5R01NS107523 (JKH)

National Multiple Sclerosis Society, JF-1806-31381 (JKH)

## Author contributions

Conceptualization: MDK, PMK

Data curation: PG, EA, MDS, MDK, PMK

Formal analysis: PG, EA, SS, MDS, WHG, JL, YG, MDK, PMK

Funding acquisition: JKH, PAC, MDK, PMK

Investigation: PG, EA, SS, MDS, WHG, JL, MB, JH, MFD, XD, SH

Project administration: MDK, PMK

Resources: LMH, JPA, JKH, PAC, MDK, PMK

Supervision: LMH, JPA, JKH, PAC, MDK, PMK

Visualization: PG, EA, SS, MDS, WHG, JL, MDK, PMK

Writing – original draft: MDK, PMK

Writing – review & editing: PG, EA, SS, MDS, WHG, JL, MB, JH, MFD, JKH, PAC, MDK, PMK

## Data and materials availability

All data are available in the main text or the supplementary materials.

