## Supplementary Materials for "PKC modulator bryostatin-1 therapeutically targets CNS innate immunity to attenuate neuroinflammation and promote remyelination"

### Materials and Methods

### Immunoblotting (Western blot)

MGCs were washed with PBS, and lysates were prepared by incubation in ice-cold RIPA buffer (50 mM Tris pH 7.4, 150 mM NaCl, 1% triton, 0.5% sodium deoxycholate, 0.1% SDS, 1 mM EDTA) supplemented with protease inhibitors. Lysates were cleared by centrifugation, and protein concentration was measured by Bradford assay. Lysates were mixed with SDS sample buffer, boiled, and resolved by SDS-PAGE. Bands were transferred to PVDF Immobilon P membranes (Millipore) using a wet transfer, blocked in TBS-Tween containing 5% milk, and probed overnight at 4°C with primary antibody against Arg1 (Santa Cruz Biotechnology, sc-271430). The membranes were then incubated for 1 hour with HRP-conjugated secondary antibody (Jackson ImmunoResearch). Immunoblots were visualized using the SuperSignal West ECL system (Thermo Fisher Scientific) followed by film exposure. Blots were then stripped using Restore Western blot stripping buffer (Thermo Fisher Scientific), blocked again as above, and probed overnight at 4°C with HRP-conjugated primary antibody against actin (Abcam), prior to visualization as above. Quantification was performed using ImageJ software.

### MGC IF staining

MGCs were plated on glass bottom 35-mm dishes and cultured in DMEM/F12 for 14 days followed by treatment. The cells washed three times with PBS and fixed with prewarmed (37°C) 4% PFA, 5 mM $MgCl_2$ (Sigma), 10 mM EGTA (Thermo Fisher Scientific), and 4% sucrose (Sigma-Aldrich) for 5-10 min at RT. The cells were then washed twice in TBS and permeabilized with 1% Triton X-100 in TBS for 5 min, followed by incubation in blocking solution (5% donkey serum, 5% goat serum in TBS) for 45 min at RT. The cells were then incubated with primary antibody rabbit anti-Arg1 (1:500) and rat anti-CDllb (1:250) in dilution buffer (20% blocking buffer in TBS) overnight at 4°C. The cells were then washed four times with the dilution buffer for 5 min. Appropriate Alexa Fluor conjugated secondary antibodies (1:500) in dilution buffer were added for 2 hours at RT and then stained with DAPI. Images were taken on Zeiss LSM 700 Confocal. Quantification was performed using ImageJ software.

**Figures**

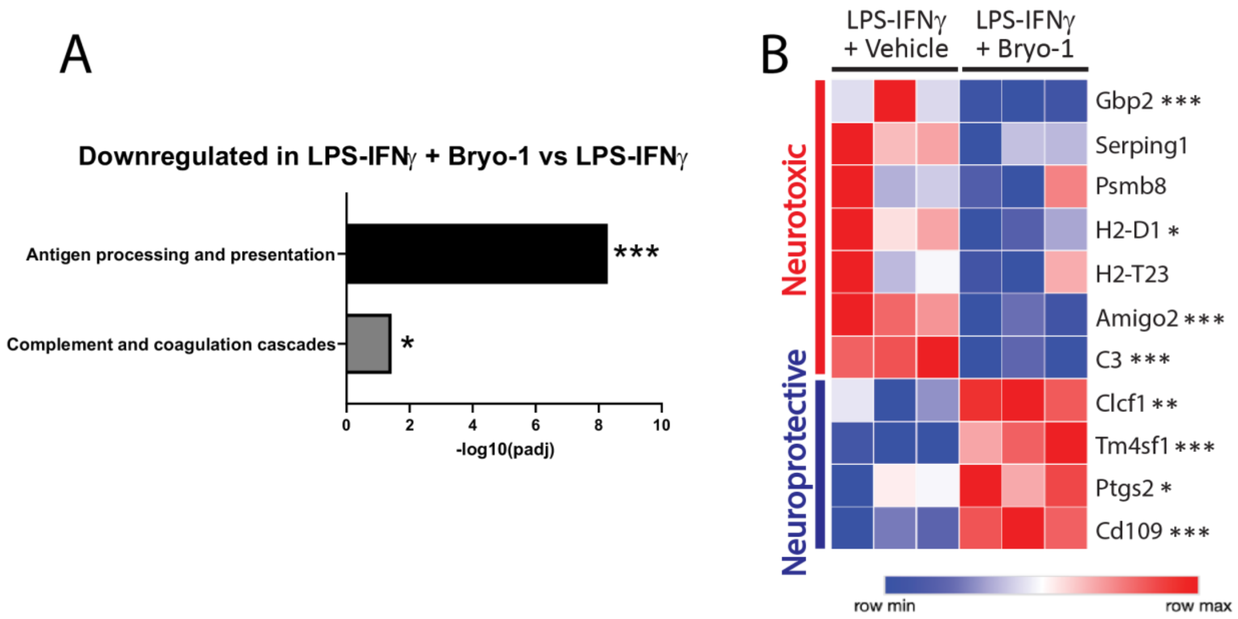


**Fig. S1. Bryo-1 inhibits pro-inflammatory responses in MGCs stimulated with LPS and IFNγ**. Mouse cortical MGCs were treated for 24 hours with LPS (100 ng/ml) and IFNγ (20 ng/ml) plus either vehicle or bryo-1 (50 nM), followed by RNA-seq analysis. **(A)** KEGG enrichment analysis highlighting key inflammatory pathways downregulated by bryo-1 treatment. **(B)** Bryo-1 modulates inflammatory astrocyte transcriptional programs induced by LPS and IFNγ in mixed glial culture. Data derived from 3 biological replicates per condition. * $p<0.05$, ** $p<0.01$, *** $p<0.001$ using DESeq2 adjusted for false discovery rate.

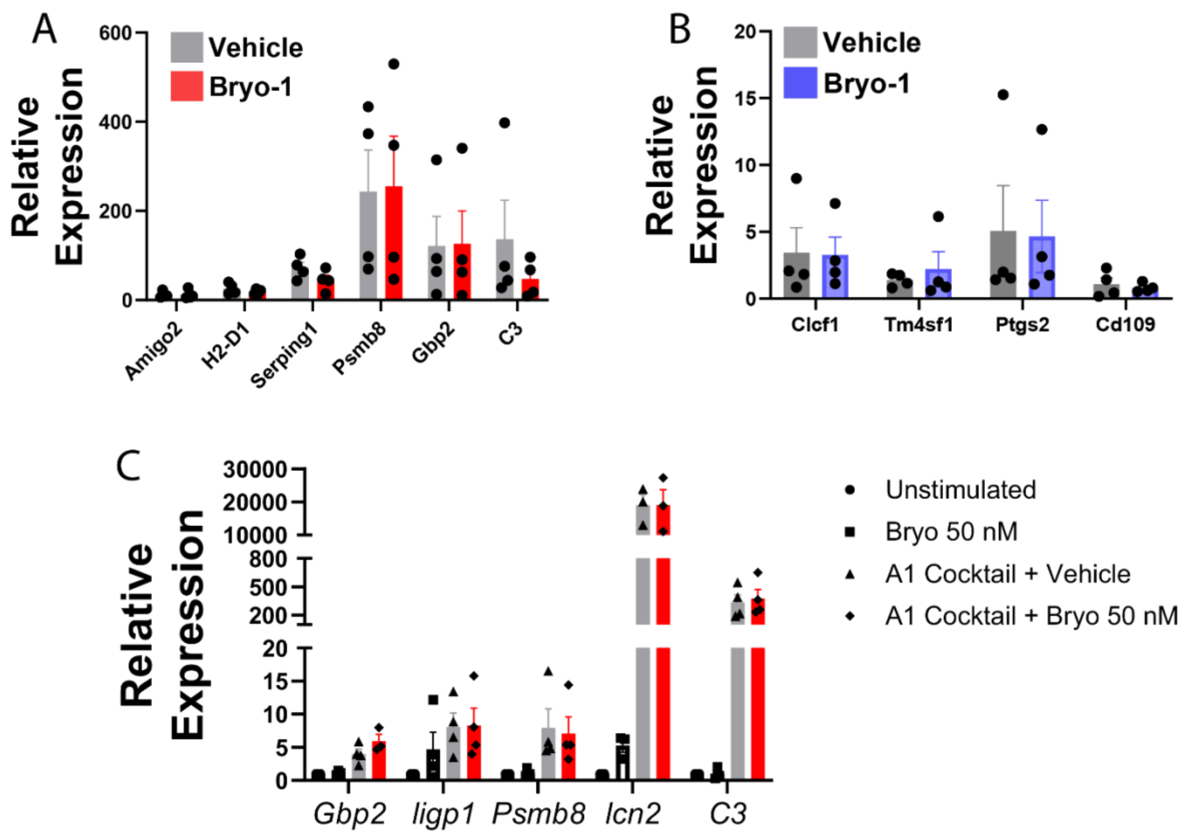


**Fig. S2. Direct treatment of astrocytes with bryo-1 has no effect on inflammatory astrocyte gene expression**. **(A, B)** Primary mouse astrocytes were treated for 24 hours with bryo-1 (50 nM) or vehicle along with conditioned media from microglia stimulated with LPS (100 ng/ml) and IFNγ (20 ng/ml), followed by qPCR for neurotoxic **(A)** and neuroprotective **(B)** astrocyte genes. **(C)** Primary mouse astrocytes were treated directly with bryo-1 (50 nM) or vehicle in the presence or absence of neurotoxic astrocyte polarization cocktail (30 ng/ml TNF, 3 ng/ml IL-1α, 400 ng/ml C1q). Expression of reactive astrocyte genes was then evaluated by qPCR. Data represent mean ± SEM from n=3-4 independent experiments.

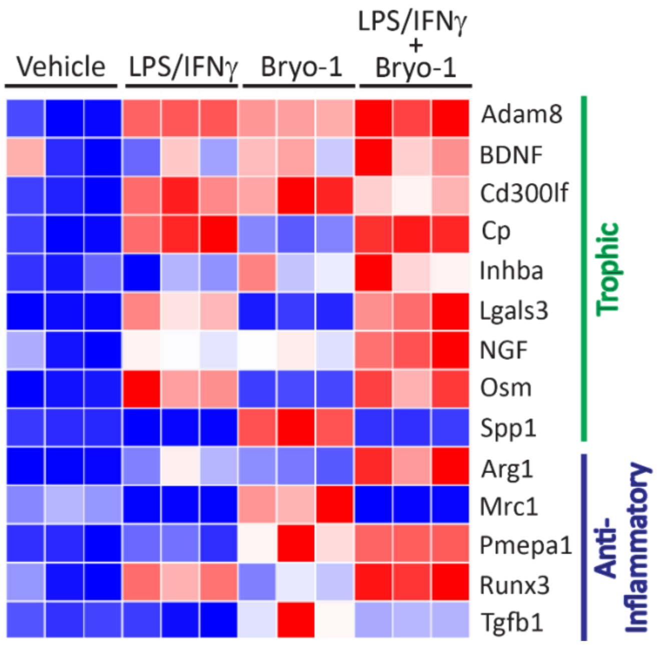


**Fig. S3. Bryo-1 promotes trophic and anti-inflammatory responses in MGCs stimulated with LPS and IFNγ.** Mouse cortical MGCs were treated for 24 hours with LPS (100 ng/ml) and IFNγ (20 ng/ml) plus either bryo-1 (50 nM) or vehicle, followed by RNA-seq analysis. Data derived from three biological replicates per condition.

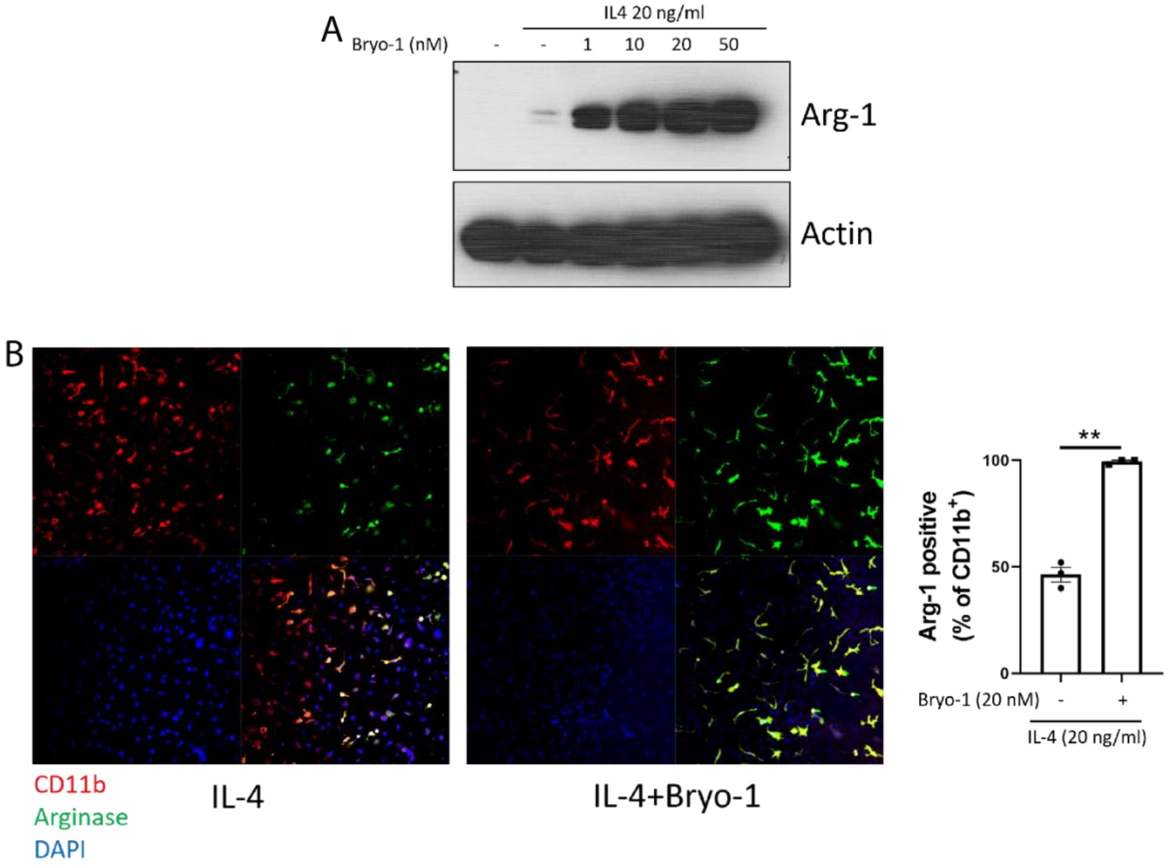


**Fig. S4. Bryo-1 augments Arg1 expression exclusively in CD11b$^+$ cells within MGCs**. Mouse cortical MGCs were treated for 24 hours with bryo-1 or vehicle ± IL-4 (20 ng/ml). **(A)** Representative immunoblot from n=3 experiments, demonstrating that bryo-1 potently augments Arg1 expression at low nanomolar doses in the presence of IL-4. Arg1 protein expression was not detectable at the protein level with bryo-1 alone (not shown). **(B, C)** Increased Arg1 expression from bryo-1 treatment (20 nM) occurs exclusively in CD11b$^+$ microglia/macrophages. **(B)** Representative images demonstrating that all Arg1$^+$ cells are also CD11b$^+$. **(C)** Quantification of Arg1 expression in CD11b$^+$ cells, as assessed by immunofluorescence. Bryo-1 induces Arg1 expression in close to 100% of CD11b$^+$ cells. Data represent mean ± SEM from n=3 experiments. ** $p < 0.01$ by two-tailed student's t-test.

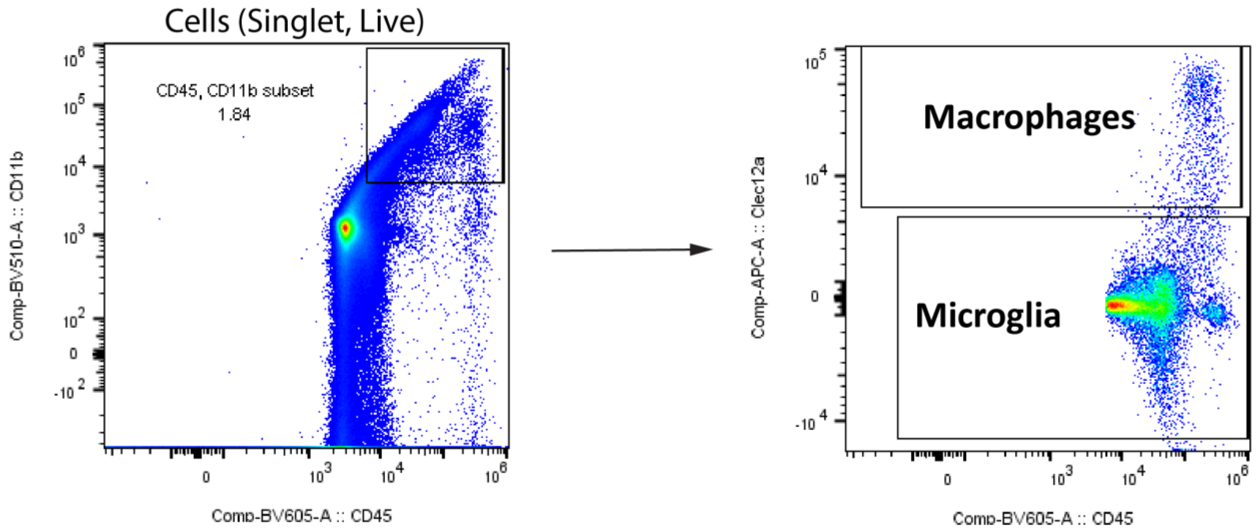


**Fig. S5. Gating strategy for microglia and macrophages applied to ex vivo phagocytosis assay shown in Fig. 3B-D**. Microglia were defined as $CD45^{+}CD11b^{+}Clec12a^{-}$, whereas macrophages were identified as $CD45^{+}CD11b^{+}Clec12a^{+}$.

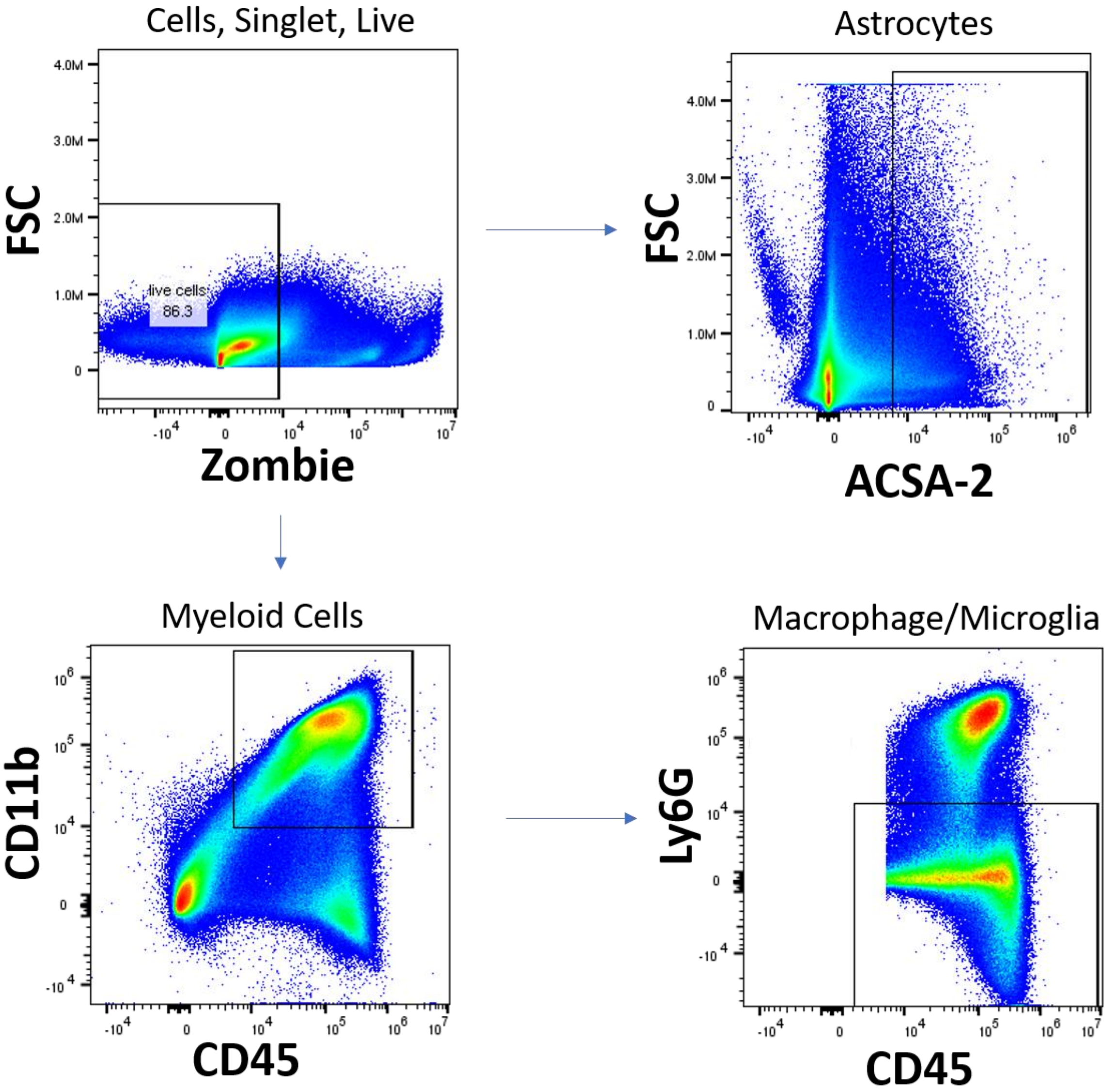


**Fig. S6. Gating strategy for spinal cord microglia/macrophages and astrocytes, corresponding to Fig. 4E and 4F**.

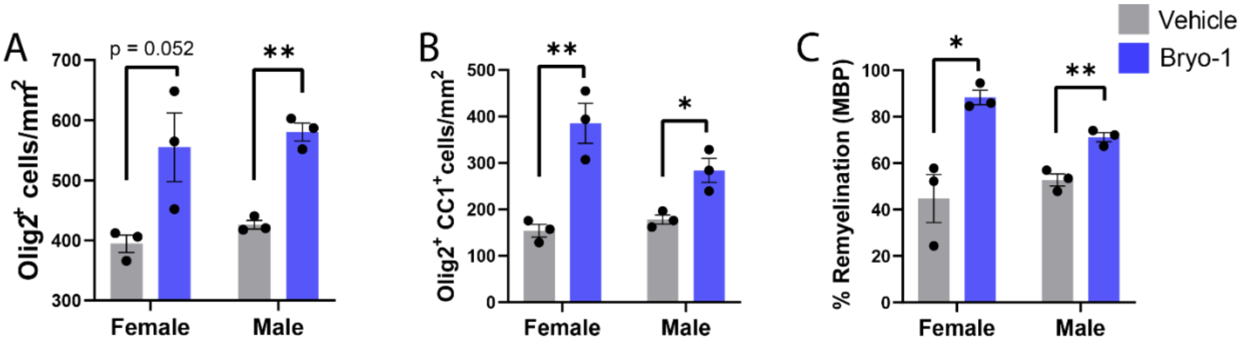


**Fig. S7. Systemic treatment with bryo-1 augments OL differentiation and remyelination in both male and female mice following focal demyelination**. Experimental design was as described in Fig. 5A. Focal demyelination was produced in the ventral spinal cord via stereotaxic injection of LPC. Beginning at 2 dpl, mice were treated with bryo-1 (35 nmol/kg) or vehicle every other day via IP injection. **(A)** Total OL-lineage cells ($Olig2^+$), **(B)** differentiating OL ($Olig2^+CC1^+$), and **(C)** remyelination (MBP expression) were then quantified at 15 dpl, as in Figure 6, with animals segregated by sex. Data represent mean ± SEM from n=3 mice per group. * $p<0.05$, ** $p<0.01$ by two-tailed student's t-test.
